# Neurophysiological evidence for cognitive map formation during sequence learning

**DOI:** 10.1101/2021.09.07.459302

**Authors:** Jennifer Stiso, Christopher W. Lynn, Ari E. Kahn, Vinitha Rangarajan, Karol P. Szymula, Ryan Archer, Andrew Revell, Joel M. Stein, Brian Litt, Kathryn A. Davis, Timothy H. Lucas, Dani S. Bassett

## Abstract

Humans deftly parse statistics from sequences. Some theories posit that humans learn these statistics by forming *cognitive maps*, or underlying representations of the latent space which links items in the sequence. Here, an item in the sequence is a node, and the probability of transitioning between two items is an edge. Sequences can then be generated from walks through the latent space, with different spaces giving rise to different sequence statistics. Individual or group differences in sequence learning can be modeled by changing the time scale over which estimates of transition probabilities are built, or in other words, by changing the amount of temporal discounting. Latent space models with temporal discounting bear a resemblance to models of navigation through Euclidean spaces. However, few explicit links have been made between predictions from Euclidean spatial navigation and neural activity during human sequence learning. Here, we use a combination of behavioral modeling and intracranial encephalography (iEEG) recordings to investigate how neural activity might support the formation of space-like cognitive maps through temporal discounting during sequence learning. Specifically, we acquire human reaction times from a sequential reaction time task, to which we fit a model that formulates the amount of temporal discounting as a single free parameter. From the parameter, we calculate each individual’s estimate of the latent space. We find that neural activity reflects these estimates mostly in the temporal lobe, including areas involved in spatial navigation. Similar to spatial navigation, we find that low dimensional representations of neural activity allow for easy separation of important features, such as modules, in the latent space. Lastly, we take advantage of the high temporal resolution of iEEG data to determine the time scale on which latent spaces are learned. We find that learning typically happens within the first 500 trials, and is modulated by the underlying latent space and the amount of temporal discounting characteristic of each participant. Ultimately, this work provides important links between behavioral models of sequence learning and neural activity during the same behavior, and contextualizes these results within a broader framework of domain general cognitive maps.

## INTRODUCTION

A diverse range of behaviors requires humans to parse complex temporal sequences of stimuli. One can study this ability by exposing individuals to sequences evincing precise statistics, and by measuring how individuals react to or remember the stimuli. Sequence statistics can be fixed by (1) an underlying graph, or latent space, defining allowable transitions between stimuli, and by (2) a walk through the graph that determines which of the allowable transitions are taken and with what frequency (**Fig. 1A**). The graph representation of the latent space brings with it a rich toolbox of methods to quantify latent space topologies that are especially well-suited for abstract relational spaces connecting discrete objects[1]. Recent studies have revealed that humans are sensitive to transition probabilities between neighboring elements[2, 3], higher-order statistical dependencies between non-neighboring elements like triplets or quadruplets[4], and the global structure of the graph[5, 6]. All of these relationships are important for naturalistic learning. For example, when learning a language, both human and artificial language processing algorithms require knowledge of which words tend to follow which others (transition probabilities), as well as about the grammar of sentences, structures of thought, and designs of paragraphs (higher-order structure)[7, 8]. Accordingly, sensitivity to these relationships predicts language ability and problem solving skills[9–11].

**FIG. 1.**
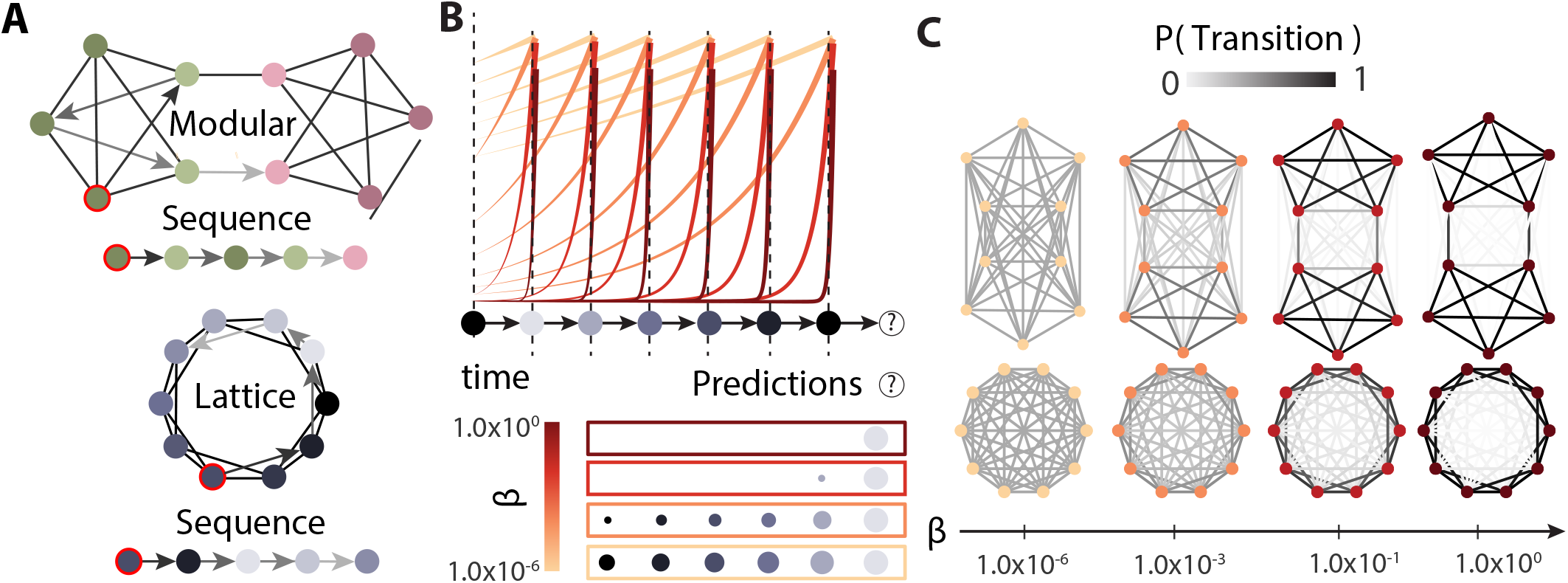
Schematic of latent space learning. *(A)* Visualization of the two graph types used to generate stimulus sequences in this study: modular (left) and lattice (right). An example sequence generated from a random walk on the graph, denoted by arrows, is shown below each visualization. *(B)* A schematic of how temporal discounting of previous stimuli leads to different predictions about which stimulus is likely to appear next in a sequence. As someone experiences each stimulus in a sequence, they will update their estimation of the latent space with the context preceding the currently viewed stimulus. The amount that each previous stimulus contributes to the context is given by the height of the colored line at the time that a given stimulus occurred. Different colors correspond to different amounts of temporal discounting or *β* values. More yellow colors indicate less temporal discounting. People can use their estimate of the latent space to predict an upcoming stimulus, indicated by a ‘(?)’. The size of each stimulus in the prediction indicates how likely that stimulus is to be the next node in the sequence. The size of a stimulus in a colored box is proportional to the height of the line of the same color when it crosses that stimulus’s presentation. We note that smaller values of *β* correspond to shallower discounting, leading to a larger range of predictions. *(C)* A visualization of how different values of *β* result in different latent space estimations. As *β* approaches 0, all transitions are estimated to be equally likely. As *β* approaches inf, estimations converge to the true structure.

Computational models of behavior that require learning an underlying latent space bear a striking resemblance to those used for learning and navigating Euclidean or abstract relational spaces[12, 13]. Moreover, similar brain regions have been implicated in all three kinds of cognitive tasks[14–16]. However, this level of generalization across task domains has been difficult to replicate in artificial intelligent systems and remains an active area of research[17, 18]. Work in sequence, relational, and spatial learning suggests that individuals may represent internal estimations of the latent spaces as cognitive maps that can be referenced during navigation and problem solving[19–22]. Recent progress in task generalizability in artificial systems has used similar techniques[17, 18]. Uncovering the processes that guide latent space estimation, and investigating how these processes are implemented in the brain, will deepen our understanding of how humans map the world around them, and provide suggestions for artificial intelligence.

Some mathematical models of latent space estimation rely on individuals building internal estimates of which stimuli in the space are likely to follow which others[23– 25]. Acquired through exploration, these estimates can be used to make predictions about which stimuli are likely to come next, and therefore allow individuals to navigate the space to reach desired goals[23]. If we were designing a system to learn latent spaces, one strategy for building estimates would be to perfectly remember and log each observed transition, and then to make predictions from that stored estimate. Although such estimates are accurate, they require the learner to store each observed transition, a requirement that is not evidenced in or expected from human behavior[13, 23, 26]. Instead, if estimates of future stimuli incorporate a broader, discounted temporal context, then some of the speed and flexibility of navigation can be restored, although at a cost to the fidelity of the estimate of the latent space[13] (**Fig. 1B**).

Temporally extended models do not recreate the exact latent space of the true environment, but their modifications can have important behavioral benefits (**Fig. 1C**). For example, artificial intelligent agents using temporal discounting can quickly navigate to rewards in new environments and flexibly respond to changes in strategies or goal locations by utilizing paths they have not explicitly traveled before[23, 25]. Without the modifications from temporal discounting providing an extended context of future paths in space resulting from a given action, agents would only be able to traverse paths they had already encountered, which would limit their flexibility. Additionally, when applied to the free recall of word sequences, these models replicate the ability of humans to remember words presented in similar contexts[27]. In these temporally extended models, when predicting which state is likely to follow the current state, the agent down-weights stimuli likely to occur far into the future relative to those in the near future, hence the term *discounting*. These temporally discounted estimates of the latent space can be constructed by applying the same discounting to the history of the previously visited stimuli[13, 25] (**Fig. 1B**). Notably, temporal discounting is a biologically feasible process and can be implemented in brain regions thought to be important for building and manipulating cognitive maps: the medial temporal and prefrontal cortices[22, 28]. Activity in medial temporal lobes has been shown to be more reflective of these discounted estimates of the latent space than the true latent space[29]. Taken together, these behavioral and neural insights support the conclusion that humans use temporally discounted estimates of latent spaces to solve a diverse set of problems.

When constructing representations of latent spaces, the brain must balance the need to accurately extract important features from the environment with the pressure to minimize resource consumption[15, 30]. This balance between compressing information and retaining important features is evidenced behaviorally in the tendency to better remember events or items that occur within a given temporal context, rather than spanning multiple contexts[31]. The medial temporal lobe is thought to facilitate the separation and generalization of contexts by identifying key features of estimated latent spaces from low dimensional projections[22]. These lower dimensional projections can serve to identify important features of the space that might be relevant for decision making, such as modules of similar items in relational spaces[32] and borders in physical spaces[22]. For cognitive maps specifically, these processes are thought to occur in the entorhinal cortex, although evidence of similar low dimensional bases in humans have been found in other regions[20, 22, 33]. Additionally, other medial temporal structures including the hippocampus have been modeled as variational autoencoders, which compress incoming sensory and structural information in order to predict future stimuli across domains[18]. Further verification that important task features can be identified from a low dimensional basis of neural activity outside of Euclidean spatial navigation would help support the generalizability of these processes. Additionally, explicit mappings between individual variations in the estimates of the latent space, and the identification of features in low dimensional space, could help us better understand the links between balancing the compression of neural activity and the need for robust behavior. Behavioral evidence suggests that all these features must be made available after relatively few exposures of different stimuli so that they can be used to make decisions[34]. Neural recordings taken during latent space learning could help clarify the timescale over which these neural features arise.

Here, we seek to better understand the neurophysiological basis of temporally discounted latent space estimation in humans. Additionally, we wish to test for similarities and divergences from processes of Euclidean spatial learning. To accomplish these goals, we will use an individual specific model of temporal discounting in patients undergoing intracranial EEG (iEEG) monitoring while completing a probabilistic serial reaction time task. In this task, participants see cues generated from a random walk on either a modular or lattice graph (**Fig. 1A**). To each individual’s reaction time data, we apply a maximum entropy model which determines the steepness of temporal discounting as parameterized by a single variable *β*[13]. This parameter also determines the structure of the corresponding estimates of the latent space for that individual. We then use representational similarity analysis to identify the electrode contacts whose activity is most similar to the estimated latent space and identify common regions involved across participants. This analysis allows us to determine whether our model’s estimation of latent spaces is reflected in neural activity, and also whether the regions involved are consistent across individuals and previously implicated in Euclidean space navigation. We find that for activity aligned to the stimulus (stimulus-locked), structures in the lateral and medial temporal lobe most often reflect the estimated latent space. In activity aligned to the response (response-locked), this similarity with the latent space shifts to frontal and premotor areas. We next tested whether low dimensional neural activity could easily identify features of the latent space, as it does in Euclidean spatial learning. We find robust separability of modules in neural activity, consistent with the identification of borders and clusters in Euclidean and relational learning. Lastly, we wish to extend our understanding of the temporal dynamics of latent space estimation. In our sample of neural data, we find that neural activity reflects the latent space within 500 stimulus exposures, and that the steepness of temporal discounting and the structure of the underlying graph influence the learning rate.

Ultimately, our study provides a direct comparison between the distinct processes of latent space learning, coupled with an evaluation of their neurophysiological underpinnings. Additionally, it provides preliminary measurements of the timescales upon which latent space estimations are formed, and an accounting of which factors influence their development. Lastly, we provide clear future directions for model development, and point out areas where neural data diverge from theoretical predictions.

## METHODS

### Participants

All participants provided informed consent as specified by the Institutional Review Board of the University of Pennsylvania, and study methods and experimental protocols were approved by the Institutional Review Board of the University of Pennsylvania.

We recruited 50 unique participants to complete our study on Amazon’s Mechanical Turk—an online marketplace for crowdsourced work. Worker identifications were used to exclude any duplicate participants. Twenty-five of the participants completed a task with a sequence generated from a modular graph, and the other 25 participants performed the same task with a sequence generated from a ring lattice graph. All participants were paid $10 for their time (≈ 20 minutes). Three individuals started, but did not complete the task, leaving the sample size at 47 individuals. Interested candidates were excluded from participating if they had completed similar tasks for the lab previously[6, 13].

#### iEEG cohort

There were a total of 13 participants (10 female, mean age 33.9 years). See Supplemental Table T1 for full demographics. This included 3 participants who completed a pilot version of the task that was largely similar. These participants were included to increase the number of participants when data collection paused during the COVID-19 pandemic. Two of these 13 participants did not have electrophysiological recordings that were synchronized with the task recordings; accordingly, these two participants were only included in behavioral analyses.

### Behavior

We test each participant’s ability to learn the structure underlying a temporal sequence of stimuli by having them perform a probabilistic motor response task using a keyboard in both the iEEG and mTurk cohorts. We will first outline elements common to both tasks here, and then highlight differences.

#### Common experimental setup and procedure

First, participants were instructed that “In a few minutes, you will see 10 squares shown on the screen. Squares will light up in red as the experiment progresses. These squares correspond with keys on your keyboard, and your job is to watch the squares and press the corresponding key when the square lights up as quickly as possible to increase your score. The experiment will take around 20 minutes”. For some participants, the sequence of stimuli was drawn from a random traversal through a modular graph (**Fig. 1A**, left); for other participants, the sequence of stimuli was drawn from a random traversal through a ring lattice graph (**Fig. 1A**,right). Both graphs have 10 distinct nodes, each of which is connected to four other nodes. Thus, the only difference between the two graphs lies in their higher-order structure. In the modular graph, the nodes are split into 2 modules of 5 nodes each, whereas in the lattice graph, the nodes are connected to their nearest and next-nearest neighbors around a ring. For each participant, the 10 stimuli are randomly assigned to the 10 different nodes in either the modular or lattice graph. The random assignment of stimuli to nodes ensures that modules are not distinguished by any stimulus features. Stimuli were each represented as a row of ten gray squares. Each square corresponds to and mimics the spatial arrangement of a key on the keyboard (**Fig. 2A**). To indicate a target key that the participant is meant to press, the corresponding squares is outlined in red (**Fig. 2B**). If an incorrect key was pressed the message “Error!” displayed on the screen until the correct key was pressed. Participants had a brief training period (10 trials) to familiarize themselves with the key presses before engaging in the task for 1000 trials, which is a sufficient number of trials for participants to learn the structure of a similarly sized modular network[6]. To ensure that participants remain motivated and engaged for the full 1000 trials, participants receive points based on their average reaction time at the end of each of 4 stages (every 250 trials). The duration of the task is determined by how quickly participants respond, but on average it takes approximately 20 minutes. On average, participants in the mTurk cohort were 94.0% ± 3.76% accurate, and participants in the iEEG cohort were 97.7% ± 2.50% accurate.

**FIG. 2.**
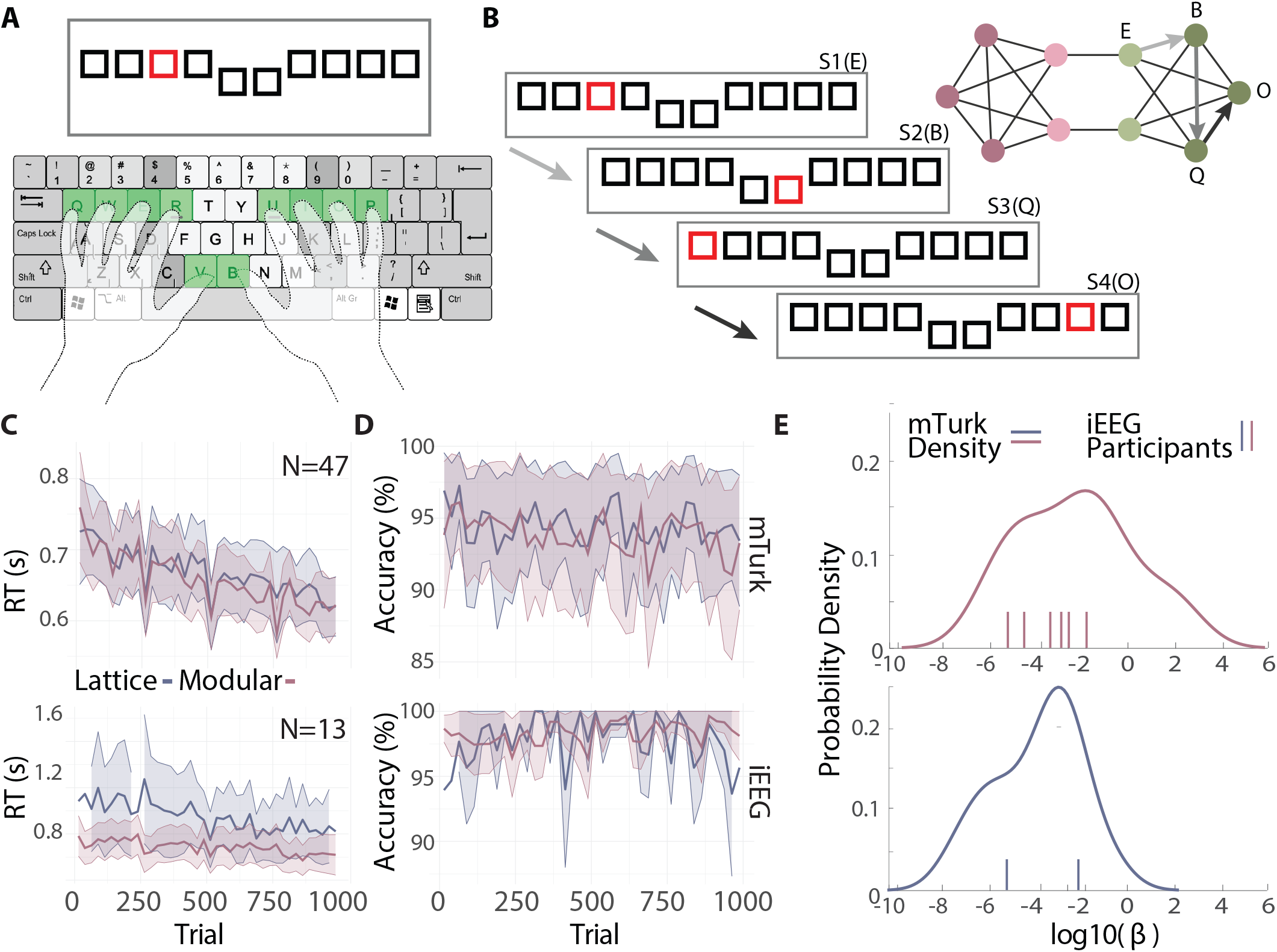
Task performance. *(A)* The hand position the participants use to complete the task. *(B)* An example of 4 trials from the experiment. The first stimulus (S1) shows the third key highlighted in red, which corresponds to the letter *E*. This key maps to the light green node in the graph to the right. After the participant presses the correct key, they will advance to next trial, which in this case is the key *B. (C)* Average reaction time over trials for the mTurk (top) and iEEG (bottom) cohorts. Reaction times are shown for both modular (pink) and lattice (blue) graphs. For visualization purposes, reaction times were averaged across participants for each trial. Those average reaction times were then binned into 25-trial bins. Shaded regions indicate the average standard error across participants for each bin. *(D)* The same plots as in panel *(C)*, but for accuracy. *(E)* Distributions of *β* values for both cohorts. Participants who saw modular graphs are shown in pink on the top; participants who saw lattice graphs are shown in blue on the bottom. Plots were separated spatially to avoid overlap between individual data points. iEEG cohort *β* values are shown with tick marks rather than as a population density due to the small sample size.

#### mTurk experiment

Because no experimenter could be present for online mTurk data collection, a few additional measures were put in place to ensure that participants understood and were engaged with the task. First, participants were given a short quiz to verify that they had read and understood the instructions before the experiment began. If any questions were answered incorrectly, participants were shown the instructions again and asked to repeat the quiz until they answered all questions correctly. Additionally, participants were instructed that if they took longer than 1 minute to respond to any given trial, the experiment would end and they might not receive payment.

#### iEEG experiment

A member of the Hospital for the University of Pennsylvania (HUP) research staff was present during the experiment to ensure that participants understood the instructions. De-identified demographic information was collected and shared from all participants as part of the HUP research protocol. This information included age, race, and sex assigned at birth, as well as an estimate of how much of their day the participant typically spent typing at a computer. The iEEG experiment, unlike the mTurk experiment, also needed to be synchronized to ongoing neural recordings. To synchronize task events with neural recordings, the iEEG participants completed the task with a photodiode attached to the laptop where the test was being administered. A white square would appear in the lower corner of the screen when a stimulus appeared on the screen, which would be replaced by a black background when the correct response was made. The photodiode would record these luminance changes on the same system that was recording neural data, so that the two could be synchronized.

Participants in the cohort were also given the option to complete a second session of the same experiment with the same graph the following day. This option was taken by 2 participants. Because data collection was interrupted by the global pandemic, we also include three pilot iEEG participants who completed an earlier version of the task that did not contain breaks or points, but was otherwise identical.

#### Linear mixed-effects models

We used linear mixed-effects models to test whether each participant’s reaction time decreased with increasing trial number. We took this decrease in reaction time as evidence that participants were learning the probabilistic motor response task. Before fitting the mixed-effects models, we excluded trials that were shorter than 50 ms, or longer than 2 standard deviations above that participant’s mean reaction time. Short trials were removed because 50 ms is not long enough to see and respond to a stimulus. We also excluded any incorrect trials. All participants in both cohorts had accuracy greater than 80%.

Mixed-effects models were fit using the lme4 library in R (R version 3.5.0; lme4 version 1.1-17), using the lmer() function for continuous dependent variables and the glmer() function for categorical dependent variables. Predictors were centered to reduce multicollinearity. Some models of accuracy did not converge with the full set of variables, so variables were removed via backwards selection with reaction time model *p*-values until the accuracy model converged. Due to the slight task differences between iEEG and mTurk cohorts, different models were used to test for learning in each cohort. For the mTurk cohort, the reaction time model was *reaction_time* ∼ *trial* ∗ *graph* + *stage* ∗ *graph* + *finger* + *hand* + *hand_transition* + *recency* +(1 + *trial* + *recency*|*participant*). The accuracy model was *correct* = *trial* ∗*graph*+*stage*∗*graph*+*finger* +*hand_transition*+ *recency* + (1 + *trial*|*participant*). Here, *hand_transition* indicated whether the current trial used a different hand than the previous trial, and *stage* indicates the set of 250 trials, ranging from 1 to 4. For the iEEG cohort, the reaction times model was *reaction_time* ∼ *trial*∗*graph* + *stage*∗*graph* + *sex* + *age* + *typing_skill* + *finger* + *hand* + *hand_transition* + *session* + *points* + *recency* + (1 + *trial* + *recency*|*participant*). The model for accuracy was *correct* ∼ *squared_trial*∗*graph*+*stage*∗*graph* + *finger* + *hand_transition* + *session* + *recency* + (1 + *squared*_*t*_*rial*|*participant*). Here, *session* indicated whether the data were taken from the first or second recording session, *points* indicated whether these participants were given points according to their reaction time at breaks, and *typing_skill* was a self-reported value of how much time participants spent typing on a computer in a typical day, scaled to range from 1-4.

The *recency* term is meant to account for changes to reaction time based on the local properties of the current sequence. Participants will tend to react more quickly to items they have seen more recently[35]. To control for this effect, we included the log transform of the number of trials since the current stimulus was last seen—or the *recency*—as a covariate. The maximum number of trials was 10. This particular covariate was found to explain more variance in reaction time than other similar covariates in this data set, as well as a similar dataset collected from Ref. [6] (see **Fig S1**). The specific covariates tested were the number of times the current stimulus was last seen (not log transformed, and not capped) (*χ*^2^ test *χ*^2^ = 2448, *p <* 2.2 × 10^−16^) and the number of times this stimulus appeared in the last 10 trials (*χ*^2^ test *χ*^2^ = 1295.8, *p <* 2.2 × 10^−16^).

#### Maximum entropy model:β and Â

To estimate the amount of temporal discounting employed by each participant, we fit a maximum entropy model to the residuals of the linear mixed-effects models specified above. The model starts with the assumption that the fastest reaction times on this task would arise from accurate mental representations of the latent space. This would allow participants to accurately predict which stimuli could possibly follow any current stimulus, allowing them to react quickly to all transitions. However, these representations are costly to create and maintain because they require perfect memory of the sequence of stimuli. Allowing some inaccuracies in the memory of previous stimuli simplifies the learning process, but at the cost of erroneous predictions about future stimuli. In this model, an exponentially decaying memory distribution determines the time scale of errors in memory. The exponential form results in the fact that mistakes in memory will be temporally discounted—more likely to occur between stimuli that are temporally close than those that are temporally distant. The steepness of this discounting, and therefore the balance of cost and accuracy, is determined by a single parameter *β* that was fitted to the residuals of each participant’s reaction times. Larger *β* values result in more temporal discounting in the memory distribution, indicating that participants were less likely to make memory errors, and the errors that were made tended to occur between stimuli in close temporal proximity. By contrast, smaller *β* values would result in less temporal discounting, indicating that participants made longer range errors in their estimates of the transition graphs. Mathematically, this is achieved by defining an individual’s estimation of the latent space as *Â* = (1 – *e*^−*β*^)*A*(*I* – *e*^−*β*^*A*)^−1^, where *A* is the true latent space that defines transition probabilities between stimuli. We will explain how *β* is calculated from reaction times below.

Given an observed sequence of nodes *x*_1,…_ *x*_*t-1*_, and given a parameter *β*, our model predicts each participant’s internal estimates of transition probabilities *Â*_*ij*_(*t-*1), where *i* and *j* are different stimuli. Given a current stimulus *x*_*t*−1_, we then model the participant’s anticipation, or expectation, of the subsequent node *x*_*t*_ by 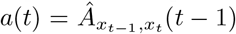. In order to quantitatively describe the reactions of a participant, we related the expectations *a*(*t*) to predictions about a participant’s reaction times 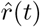, and then learned the model parameters that best fit that participant’s reaction times. The simplest possible prediction was given by the linear relation 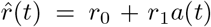, where the intercept *r*_0_ represents a participant’s reaction time with zero anticipation and where the slope *r*_1_ quantifies the strength with which a participant’s reaction times depend on their internal expectations. In total, our predictions *r*(*t*) contain three parameters (*β, r*_0_, and *r*_1_), which must be estimated from the data for each participant. To estimate the model parameters that best describe a participant’s reaction times *r*(*t*) (more specifically, their reaction time residuals from the linear mixed-effects model described above), we minimized the root mean squared prediction error (RMSE) with respect to each participant’s observed reaction times, 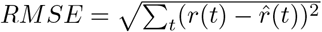. We note that, for a given *β*, the parameters *r*_0_ and *r*_1_ can be calculated using linear regression. Thus, the problem of estimating the model parameters can be restated with only one parameter; that is, by minimizing the RMSE with respect to *β*.

Because we wished to compare results from these models to neural data, we only run this analysis on each of the participants with neural data, and exclude trials that contained interictal epileptiform discharges (IEDs). To minimize the *RMSE* with respect to *β*, we began by calculating the *RMSE* along 100 logarithmically spaced values for *β* between 10^−4^ and 10. Then, starting at the minimum value of this search, we performed gradient descent until the gradient fell below an absolute value of 10^−6^. The search also terminated if *β* reached 0, or was trending towards ∞ (greater than 1000). The *β* values that were terminated at 0 or 1000 are referred to as extreme values throughout the manuscript.

Once *β* values were fitted for each participant, the estimated latent space *Â* could be obtained with the equation: *Â* = (1 – *e*^−*β*^)*A*(*I* – *e*^−*β*^*A*)^−1^, where *A* is the true latent space that defines transition probabilities between stimuli. This analytic prediction reflects the estimated latent space for a participant that viewed an infinite random walk, and does not take into account the statistics of the particular sequence observed by a given participant.

In addition to calculating each participant’s estimated latent space, we also wished to understand how the estimate would evolve over time assuming a static *β*. A participant’s expected likelihood of a transition between two elements *i* and *j* at time *t* is given by 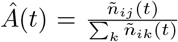, where *ñ*_*ij*_ is a participant’s recollection of the number of times they have observed stimulus *i* transition to stimulus *j*. We can then use *β* to solve for the expected number of transitions as 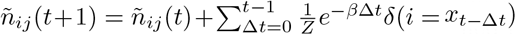. Here, *δ*(*·*) is a delta function that gives a value of 1 when its argument is true and 0 otherwise, and *Z* is a normalization constant.

### Intracranial Recordings

All patients included in this study gave written informed consent in accord with the University of Pennsylvania Institutional Review Board for inclusion in this study. De-identified data were retrieved from the online International Epilepsy Electrophysiology Portal[36]. All data were collected at a 512 Hz sampling rate.

#### Preprocessing

Electric line noise and its harmonics at 60, 120, and 180 Hz were filtered out using a zero phase distortion fourth order stop-band Butterworth filter with a 1 Hz width. This filter was implemented using the butter() and filtfilt() functions in MATLAB. For impulse and step responses of this filter, see Supplemental **Fig. S2**.

We then sought to remove individual channels that were noisy, or had poor recording quality. We first rejected channels using both the notes provided and automated methods. After removing channels marked as low quality in the notes, we further marked electrodes that had (1) a line length greater than three times the mean[37], (2) a *z*-scored kurtosis greater than 1.5[38], or (3) a *z*-scored power-spectral density dissimilarity measure greater than 1.5[38]. The dissimilarity measure was the average of one minus the Spearman’s rank correlation with all channels. These automated methods should remove channels with excessive high frequency noise, electrode drift, and line noise, respectively. All contacts selected for removal were visually inspected by a researcher with 6 years of experience working with iEEG data (JS). The final set of contacts was also visually inspected to ensure that the remaining contacts had good quality recordings by the same researcher. Including removal of contacts outside of the brain, on average, 48.87% ± 22.50% of contacts were removed, leaving 89 ± 30 contacts.

Data were then demeaned and detrended. Detrending was used instead of a high-pass filter to avoid inducing filter artifacts[39]. Channels were then grouped by grid or depth electrode, and common average referenced within each group. Recordings from white matter regions have sometimes been used as reference channels[40]. However, work showing that channels in white matter contain unique information independent from nearby grey matter motivated us to include them in the common average reference[41]. Following the common average reference, plots of raw data and power spectral densities were visually inspected by the same expert researcher with 6 years of experience working with electrocorticography data (JS) to ensure that data were of acceptable quality.

Next, data were segmented into trials. A trial consisted of the time that a given stimulus was on the screen before a response occurred. iEEG recordings were matched to task events through the use of a photodiode during task completion (see *iEEG experiment* section). Periods of high light content were automatically detected using a custom MATLAB script. Identified events were then visually inspected for quality. The times of photodiode change were then selected as the onset and offset of each trial. Two participants had poor quality photodiode data that could not be segmented, and these participants were accordingly not included in electrophysiological analyses, leaving 11 remaining participants.

Lastly, trials were rejected if they contained interictal epileptiform discharges (IEDs). IEDs have been shown to change task performance[37] and aspects of neural activity outside of the locus of IEDs[42, 43]. We chose to use an IED detector from Ref. [44] because it is sensitive, fast, and requires relatively little data per participant. This Hilbert-based method dynamically models background activity and detects outliers from that background. Specifically, the algorithm first downsamples the data to 200 Hz, and applies a 10-60 Hz bandpass filter. The envelope of the signal is then obtained by taking the square of this Hilbert-transformed signal. In 5 second windows with an overlap of 4 seconds, a threshold *k* is calculated as the mode plus the median and used to identify IEDs. The initial *k* value is set to 3.65, which was determined through cross-validation in Ref. [44]. In order to remove false positives potentially caused by artifacts, we apply a spatial filter to the identified IEDs. Specifically, we remove IEDs that are not present in a 50 ms window of IEDs in at least 3 other channels. The 50 ms window was consistent with that used in other papers investigating the biophysical properties of chains of IEDs, which tended to last less than 50 ms[45].

#### Contact localization

Broadly, contact localization followed methodology similar to Ref. [46]. All contact localizations were verified by a board-certified neuroradiologist (JMS). Electrode coordinates in participant T1w space were assigned to an atlas region of interest and also registered in participant T1w space. Brain region assignments were assigned first based on the AAL-116[47] atlas. This atlas extends slightly into the white matter directly below grey matter, but will exclude contacts in deeper white matter structures. For a list of the number of contacts in each region of this atlas, see Table T2. To provide locations for contacts outside the AAL atlas, we use the Talairach atlas[48]. Assignment of contacts to a hemisphere was also done using the Talairach atlas label. For a list of the number of contacts outside the AAL atlas in each Talairach region, see Table T3. If the contact was outside of the Talaraich atlas, then the AAL atlas hemisphere was used. If a contact was outside both atlases, then the contact name taken from iEEG.org was used (contact names include the hemisphere, electrode label, and contact label).

#### Similarity analysis

In this work, we sought to identify which electrode contacts have neural activity that reflected a participant’s estimate of the latent space in a data driven manner. To identify these contacts, we used a similarity analysis that compared *Â*, the participant’s estimation of latent space, to the similarity of neural activity evoked by each stimulus. This approach was used to abstract similarity patterns in high-dimensional neural activity into dissimilarity matrices, and allowed us to answer the question “Where does neural activity reflect the latent space?” [49]. These matrices can then be compared with similarity patterns obtained from our computational model, *Â*.

Here, we chose the cross-validated Euclidean distance as our neural similarity metric because it was shown to lead to more reliable classification accuracy when compared to other dissimilarity metrics[50]. To compute similarity matrices for each contact, we first truncated all trials of preprocessed iEEG recordings to be the same length as the trial with the shortest reaction time. If the shortest reaction time was less than 200 ms, we instead used 200 ms as the minimum length and discarded trials shorter than that. This truncation was done in three ways: (1) stimulus aligned, where the end of trials was truncated; (2) middle aligned, where the middle of trials was truncated; and (3) response aligned, where the beginning of trials was truncated. We then calculated the leave-one-out cross-validated Euclidean distance between activity evoked from each of the 10 unique stimuli. This procedure resulted in one dissimilarity matrix for each contact. To compare these matrices to the estimated latent space, we then calculated the correlation between the lower diagonal of the neural dissimilarity matrix and *Â*. Because *Â* reflects similarity rather than dissimilarity, we then multiplied the resulting correlation by -1.

To identify electrode contacts with high similarity to the latent space, we compared (1) the correlations between the neural dissimilarity and the estimated latent spaces to (2) correlations between a distribution of 100 null neural dissimilarity matrices and estimated latent spaces. Null matrices were calculated from permuted data created by first selecting a random trial number and then splitting and reversing the order of trials at that point. For example, if 128 were drawn as a random trial number, the corresponding permuted dataset would be neural data from trials 129 − 1000 followed by neural data from trials 1 − 128 matched with stimuli labels from the correct order of trials (1 − 1000). This model preserved natural features of autocorrelation in the neural data, unlike trial shuffling models[51]. Contacts were determined to have activity similar to the latent space if they met two criteria: (1) the correlation between the neural dissimilarity matrix and the *Â* was greater than at least 95 null models, and (2) the correlation between the neural dissimilarity matrix and the *Â* was greater than the correlation between the neural dissimilarity matrix and the exact latent space *A*.

To test the specificity of our findings, we also examined the correlation between the dissimilarity matrices and a similarity space related to the lower-level features of the stimuli. We calculated a spatial similarity matrix that reflected the physical distance between stimuli on the screen. Since each stimulus consists of a single red square among 9 black squares on the screen, we calculated the Euclidean distance between each square, and used this matrix as an estimate of spatial similarity. We then repeated the process detailed above for obtaining correlations relative to permuted neural data.

#### Low dimensional projections and linear discriminability

For visualization purposes, we sought to obtain low dimensional representations of the neural dissimilarity matrices. Classical multidimensional scaling (MDS) obtains low dimensional (here, 2 dimensions) representations of Euclidean distance dissimilarity matrices that seek to preserve the distances of the original higher-dimensional data[52]. Classical multidimensional scaling was implemented using the cmds() function in MATLAB. For neural data, we first calculated a single neural dissimilarity matrix, rather than a single matrix per contact. This calculation was done by concatenating activity from every contact whose activity was similar to the latent space (see *Similarity analysis* section), and then by repeating the process outlined above.

For some analyses, we wished to compare the low dimensional representations of neural dissimilarity matrices with the low dimensional representations of estimated latent spaces. Since estimated latent spaces are not Euclidean distance matrices, classical MDS is not an appropriate dimensionality reduction technique[52]. Instead, we use principal components analysis (PCA). PCA yields the same low dimensional embedding as classical MDS when the high dimensional data are Euclidean distances, but not otherwise. We computed the principal components of the neural dissimilarity matrices and estimated latent spaces in MATLAB using the pca() function. The scaled and centered data were then projected onto the first 2 principal components to obtain 2 coordinates for each node.

From these low dimensional data, we next sought to assess estimates of discriminability between modules. Module discriminability was calculated as the loss from a linear discriminant analysis. A linear classification model was fit to the low dimensional coordinates using the fitdiscr() function in MATLAB. The proportion of nodes that were incorrectly classified using the best linear boundary, or the loss, was then reported as an estimate of the linear discriminability of modules.

### Statistical Analyses

Linear mixed-effects models were used to analyze reaction time data, and the results are displayed in **Fig. 2**. Mixed-effects models were used to account for the fact that trials completed by the same participant constitute repeated measures and are not independent. The estimated *β* values were evaluated with *t*-tests, and appear to be approximately normally distributed. Extreme values of *β* (0 or 1000) were removed from any statistical tests to ensure normality (see **Figs. 1** and **5**). Linear mixed-effects models were used to analyze changes in neural similarity over time, with participant included as a random effect (**Fig. 5**). A paired *t*-test was used to analyze changes in loss from a linear classifier (**Fig. 6**).

### Data and Code

Code is available in the github repository github.com/jastiso/stistical_learning. Electrophysiological data will be made available upon request from the IEEG Portal.

## RESULTS

### Quantification of learning and temporal discounting

In this work, we are interested in the neural underpinnings of latent space estimation. Before investigating the neural dynamics directly, we tested whether participants learned the latent space and responded both faster and more accurately to stimuli over time. Our cohort of interest, the iEEG cohort, were all undergoing monitoring for medically refractory epilepsy. Due to the rarity of this population, it is often difficult to get large cohorts suitable for good estimates of behavioral effect sizes. Additionally, the epileptic population in the iEEG cohort has been shown to have cognitive impairments[53], which requires tasks that have been designed to be comparatively easy and quick to complete. Due to these challenges, we also collected data from 50 participants from Amazon’s Mechanical Turk (mTurk).

In both cohorts, we were interested in the change over time of two estimates of learning: accuracy and reaction time. Across participants, we found that the average accuracy for the mTurk cohort was 94.0% ± 3.76%, with a median reaction time of 602.5 ± 134.0*ms*. For the iEEG cohort, the mean accuracy was 97.7% ± 2.50% with a median reaction time of 721.2 ± 180.9*ms*. We were also interested in determining whether the rate of learning differed between two graph types (**Fig. 1A**). We used linear mixed-effects models to assess learning based on increases in accuracy and decreases in reaction times on two time scales. The first, shorter timescale is that of individual trials; to examine learning on this timescale we tested for decreases in reaction time associated with increasing trial number. Since this task provided breaks at 250 trial stages, we also assessed learning at the longer timescale of individual stages. To examine learning on this timescale, we tested for decreases in reaction time with increasing stages. In the mTurk cohort (*n* = 47), we found that reaction times tend to decrease only at the trial level (linear mixed-effects model *F*_*trial*_ = 16.1, *p*_*trial*_ = 9.51 × 10^−5^, *F*_*stage*_ = 0.005, *p*_*stage*_ = 0.946; **Fig. 2C**). In the iEEG cohort (*n* = 13), we found that reaction times decrease only at the stage level (linear mixed-effects model *F*_*trial*_ = 1.16, *p*_*trial*_ = 0.320, *F*_*stage*_ = 3.86, *p*_*stage*_ = 0.049; **Fig. 2C**). For accuracy, we found that the mTurk cohort shows a significant decrease in accuracy with trials (linear mixed-effects model *z*_*trial*_ = −2.48, *p*_*trial*_ = 0.013, *z*_*stage*_ = 1.93, *p*_*stage*_ = 0.054; **Fig. 2D**). For the iEEG cohort we observe no significant linear change with trial (linear mixed-effects model *z*_*trial*_ = −0.025, *p*_*trial*_ = 0.98, *z*_*stage*_ = 0.289, *p*_*stage*_ = 0.773; **Fig. 2D**). However, we qualitatively observed a quadratic relationship, where accuracy initially increased before decreasing with trial number. We tested the statistical significance of this observation with a mixed-effects model that relates accuracy to *trial*^2^. We found that the quadratic trial estimate is a significant predictor of accuracy (linear mixed-effects model 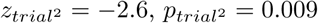; **Fig. 2D**).

We also sought to determine whether these effects differed by graph type. In the mTurk cohort, we found that there was no difference in reaction time (linear mixed-effects model *F*_*graph*_ = 0.013, *p*_*graph*_ = 0.910) or in learning rate (*F*_*trial*\**graph*_ = 0.043 *p*_*trial*\**graph*_ = 0.834, *F*_*stage*\**graph*_ = 0.002, *p*_*stage*\**graph*_ = 0.966; **Fig. 2C**) between the graphs. There were also no significant changes in accuracy associated with graph type (linear mixed-effects model *z*_*graph*_ = −0.186, *p*_*graph*_ = 0.853, *z*_*trial*\**graph*_ = −0.818, *p*_*trial*\**graph*_ = 0.414, *z*_*stage*\**graph*_ = 1.121, *p*_*graph*\**stage*_ = 0.225; **Fig. 2D**). In the iEEG cohort, we found no differences in reaction time (*F*_*graph*_ = 1.63, *p*_*graph*_ = 0.300), but there was a significant interaction between learning rate and graph type at the stage level (*F*_*graph*\**trial*_ = 4.70, *p*_*trial*\**graph*_ = 0.072, *F*_*stage*\**graph*_ = 14.3, *p*_*stage*\**graph*_ = 1.52 × 10^−4^; **Fig. 2C**). There was also a significant interaction between accuracy and graph type (linear mixed-effects model, *z*_*graph*_ = −2.6, *p*_*graph*_ = 0.711, *z*_*trial*\**graph*_ = 2.30, *p*_*trial*\**graph*_ = 0.022, *z*_*stage*\**graph*_ = 1.94, *p*_*stage*\**graph*_ = 0.052; **Fig. 2D**). Overall, we found that the iEEG cohort showed evidence of learning in both accuracy and reaction time. While the mTurk cohort showed quicker decreases in reaction time, these were coupled with decreases in accuracy. Additionally, this analysis supported steeper learning on the lattice graph within the iEEG cohort, although it is important to note that only 4 participants were exposed to lattice graphs.

After we confirmed that participants learned the task, we quantified each participant’s steepness of temporal discounting. For both cohorts, we calculated the parameter *β* by fitting a maximum entropy model to the residuals of reaction times from the linear mixed-effect model discussed above. This parameter indicates the prioritization of accurate latent space estimations against the cost of those accurate representations, as evidenced by each participant’s behavior (**Fig 2A**). The parameter *β* was fit with gradient descent, assuring that the fit for each participant was comparable, with the exception of the extremes of the distribution of possible *β* values (*β* = 0 and *β* = ∞). A fitted value of *β* = 0 indicated that there is no evidence of temporal discounting in a participant’s behavior, and the corresponding estimate *Â* of the latent transition probabilities would show equally likely transitions between all nodes. A fitted value of *β* = ∞ indicated no influence of the cost of building accurate representations, resulting in an *Â* that converges to the true latent space. Because the gradient descent algorithm terminated if *β* approached 0 or ∞, we assessed the similarity of temporal discounting—operationalized as similar *β* values—between cohorts with 2 measures: (1) the percent of participants where *β* approached one of these extremes; and (2) the distribution of *β* values found between these two extremes. Additionally, all parametric statistical tests that used *β* values were applied after extreme values were removed, thus ensuring the normality of the *β* distribution.

We first examined the percentage of participants who had *β* values at the extremes of the distribution. In the mTurk cohort, we found that 55.3% (or 27 participants) had extreme *β* values. Thirteen of these values were from participants with lattice graphs, and thirteen were from participants with modular graphs. For the iEEG cohort, we found that 27.3% (or 3; 2 lattice, 1 modular) of participants had extreme *β* values. We next assessed differences in the distribution of *β* values in both cohorts. The mTurk cohort had a mean *β* value of 0.94, and showed no differences across graph type (permutation test: *p* = 0.11) (**Fig. 2E**). The iEEG cohort had a mean *β* value of 0.17 and was not statistically different from the mTurk cohort (permutation test: *p* = 0.53) (**Fig. 2E**). As these data indicate, we found similar temporal discounting levels amongst both groups, although the mTurk cohort had more participants with extreme values. We note that *β* values tend to be less than 1, indicating a high prioritization of the costs of building accurate representations. Since this amount of temporal discounting resulted in estimated latent spaces that are different from true latent spaces, we next investigated neural activity reflecting these estimated latent spaces.

### Anatomical areas where activity reflects latent space estimation

We used a similarity analysis in a data driven manner to identify which contacts showed activity with a similar structure to the estimated latent space. First, we calculated the similarity structure of neural activity by calculating the cross-validated Euclidean distance between the activity evoked for each stimulus (**Fig. 3A**). To ensure that all stimuli had activity of the same length, the last time points of all trials were removed to create epochs the length of the shortest trial. We also report results based on removing the first and middle time points to reach the same length. We then selected the contacts where this neural similarity structure was closest to the estimated latent space. These contacts were selected by two criteria (see Methods): (1) the correlation with the estimated latent space must be larger than correlations from 95 out of 100 null models; and (2) the correlation with the estimated latent space must be larger than the correlation to the exact latent space. The null matrices were calculated from neural data where the trial order had been split and reordered. We wished to compare the contacts selected for similarity to the latent space with contacts that showed similarity to other task-relevant features that were not selected from behavior. Thus, we repeated the null model comparison described above, but compared neural dissimilarity matrices to a matrix of Euclidean distances between stimuli on the screen, rather than the estimated latent space (**Fig. 3A**). We refer to this distance as the visual distance. Because we only require that contacts have similarity values greater than 95% of null models and the correlation with the exact latent space, we expect that a rate of false positives amongst contacts of close to (but less than) 5%. Therefore, we focus our discussion on regions where greater than 5% of the total contacts were retained.

**FIG. 3.**
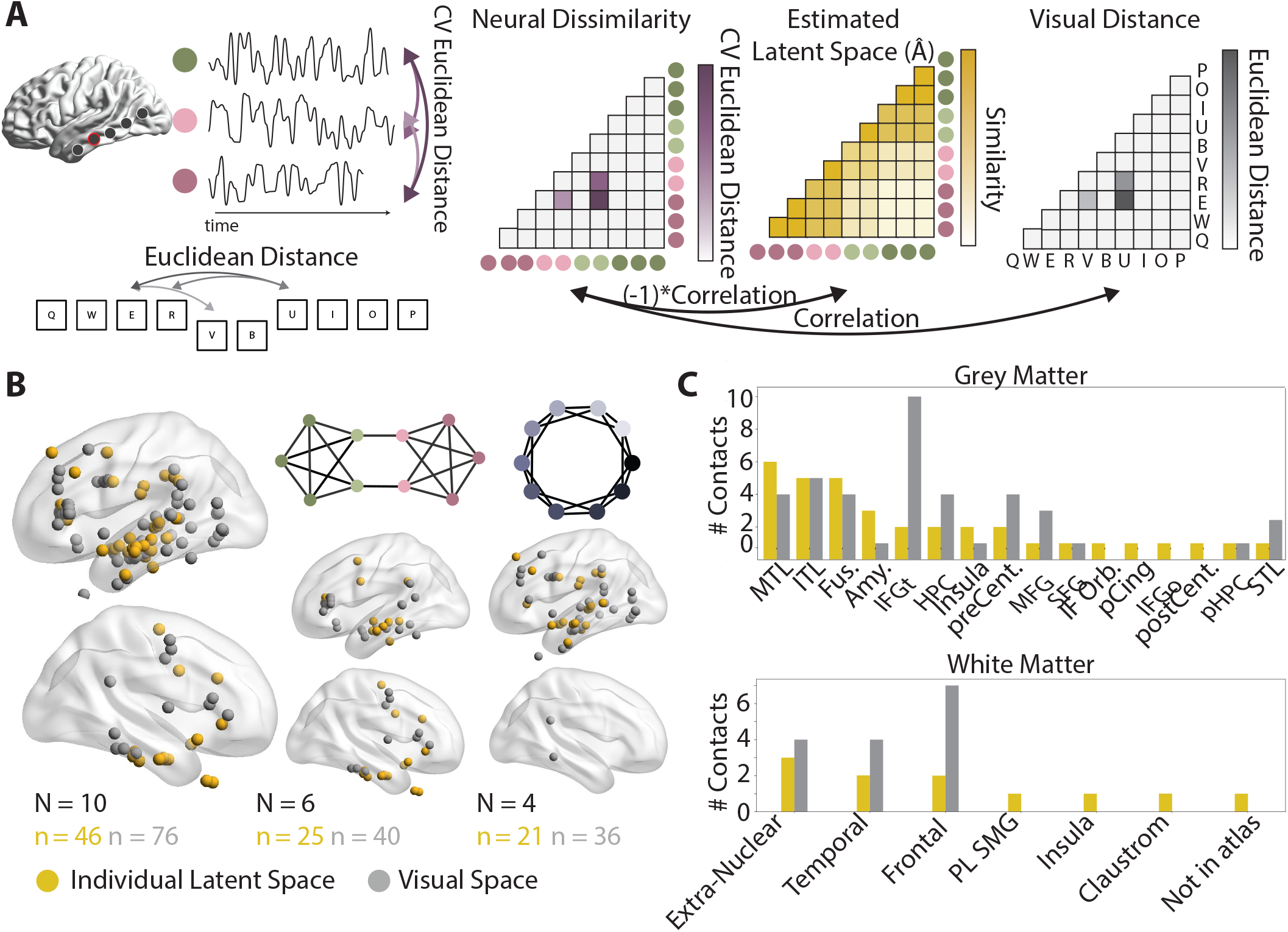
Location of contacts whose activity reflects the estimated latent space. *(A)* A schematic of the representational similarity analysis used here. First Euclidean distances (top, left) are calculated from neural responses to each stimulus to create a neural dissimilarity matrix (leftmost matrix). The Euclidean distance between highlighted stimuli are also calculated (bottom, left) and used to make a visual distance matrix (rightmost matrix). The similarity between the neural dissimilarity matrix and both the estimated latent space *Â* and the visual space are assessed with Pearson’s correlation coefficients. Because the estimated latent space is a similarity matrix, rather than a dissimilarity matrix, the correlations are multiplied by -1. *(B)* Visualization of contacts with similar activity spaces on an MNI brain for all participants (left), only those who saw sequences from a modular graph (middle), and only those who saw sequences from a lattice graph (right). Contacts whose neural dissimilarity matrices are similar to those of the latent space are shown in gold; contacts whose neural dissimilarity matrices are similar to those of the visual space are shown in grey. The quantity *N* is the number of participants in each plot, and the quantity *n* is the number of contacts in each plot. *(C)* Anatomical localization of grey (top) and white (bottom) matter contacts. Grey matter contacts are localized with the AAL atlas, and white matter contacts are localized with the Talaraich atlas. ITL: inferior temporal lobe, MTL: middle temporal lobe, STL: superior temporal lobe, Fus: fusiform, pHPC: parahippocampal gyrus, Amy: amygdala, HPC: hippocampus, IFGt: inferior frontal gyrus (pars triangularis), IFGo: inferior frontal gyrus (pars orbitalis), MFG: middle frontal gyrus, SFG: superior frontal gyrus, preCent: precentral, postCent: postcentral, pCing: posterior cingulate, PL SMG: parietal lobe supramarginal gyrus.

The resulting contacts from all participants are visualized on a shared space (MNI; **Fig. 3B**). Between 2 and 10 contacts displayed activity whose dissimilarity matrices were similar to those of the latent space per participant (**Fig S3**). Qualitatively, we observed that contacts that reflect latent and Euclidean space appear in the frontal and temporal lobes, with some overlap between the two groups. Overall, 46 (5.0%) contacts spanning all participants were identified as reflecting the latent space, and 76 (8.3%) were identified as reflecting the Euclidean space. For the latent space, 32 (5.1%) contacts were from the right hemisphere and 14 (4.5%) contacts were from the left hemisphere (**Fig. S3**). We note that we expect to select more visual than latent space sensitive contacts because visual space correlations were not required to be larger than the correlations to the true latent space. We also show separate visualizations for participants with modular and lattice graphs, respectively (**Fig. 3B**). Qualitatively, we observe a large overlap in the identified regions between the two graph types.

We next sought to localize identified contacts in each participant’s native space. The most common AAL atlas labels for latent space contacts are shown in **Fig. 3C**. We found the most common regions identified were the middle temporal lobe (6 contacts 5.0%), fusiform gyrus (5 contacts, 11.0%), inferior temporal lobe (5 contacts, 4.6%) and amygdala (3 contacts, 27.3%). The middle temporal lobe and amygdala also showed the most selectivity for the latent space compared to the visual space. Contacts located in white matter were localized with the Talaraich atlas. Most often, these contacts are in extra-nuclear, frontal or temporal sub-lobar white matter. Information for all regions is given in Supplemental **Table T3**. We note that the most common regions identified for latent space contacts were in lateral, ventral, and medial temporal lobe. While frontal contacts still showed neural dissimilarity consistent with that of the latent space, the specific anatomical location of these contacts were not consistent across participants.

The above similarity analysis was based on data that was aligned to stimulus presentation, and hence captured the initial evoked response to the stimulus. To examine whether other portions of the response data produced the same structure, we repeated this analysis with response-aligned trials (removing the first time points), and middle-aligned trials (removing the middle time points) (**Fig. S4-S5**). When trials were aligned to the response, we found that fewer contacts were identified whose activity reflected the latent space, although such contacts were still identified in each participant. Overall, we found 36 (3.9%) contacts showed activity similar to the estimated latent space, and 44 (4.8%) contacts showed activity similar to the visual space. For latent space contacts, 24 (3.9%) were in the left and 12 (4.0%) in the right hemisphere. We found one region, the fusiform gyrus (5 contacts 10.9% for response-aligned), that included multiple contacts for both response- and stimulus-aligned activity. Unlike the stimulus-aligned similarity, the response-aligned similarity also identified the inferior frontal gyrus (the pars orbitalis, pars triangularis, and pars opercularis) (4, 7.7%), insula (3, 14.3 %), and supplemental motor area (3, 60%) as important regions. For the middle-aligned activity, we found 52 (5.7%) contacts showed activity reflecting the latent space and 65 (7.1%) showed activity reflecting the visual space. Among these latent space contacts, 39 (6.3%) were in the left hemisphere and 13 (4.4%) were in the right hemisphere. We found that most identified regions overlap with those identified for stimulus- and response-aligned activity. Specifically, we found that the areas most commonly displaying activity that reflects the latent space are located in the middle temporal lobe (10, 8.3%), the fusiform gyrus (4, 8.7%), the inferior frontal gyrus (4, 7.7%), and the inferior temporal cortex (4, 3.7%). These results suggest that our findings from response- and stimulus-aligned activity are not driven by the activity of the middle of the trial. Overall, we found that early stimulus-evoked activity shows greatest similarity to the estimated latent space in higher-order temporal regions, whereas later response-locked activity shows more similarity to the estimated latent space in frontal and especially pre-motor regions.

### Module discriminability in low dimensional space

Many of our hypotheses about low dimensional projections of neural activity build upon prior evidence in the medial temporal lobe. To be more consistent with this literature, the remaining analyses considered only the stimulus-locked neural dissimilarity matrices, where the temporal lobe contacts most reflected the estimated latent space. We visualized low dimensional projections of neural activity across all of the contacts that demonstrated similarity to the estimated latent space (**Fig. 4**). These low dimensional projections were obtained for each participant by first creating a single dissimilarity matrix for all contacts whose activity was similar to the latent space, and then computing classical multidimensional scaling on those matrices. From these low dimensional projections, we can observe the diversity of estimated structures, and the ways in which they reflect and differ from the exact latent space that generated the sequences of images (**Fig. 4**). One notable property of participants who experienced sequences from modular graphs is that modules (green and pink) appear to be mostly separable. All participants appear to highly accurately separate the two modules, even when activity from diverse regions is included.

**FIG. 4.**
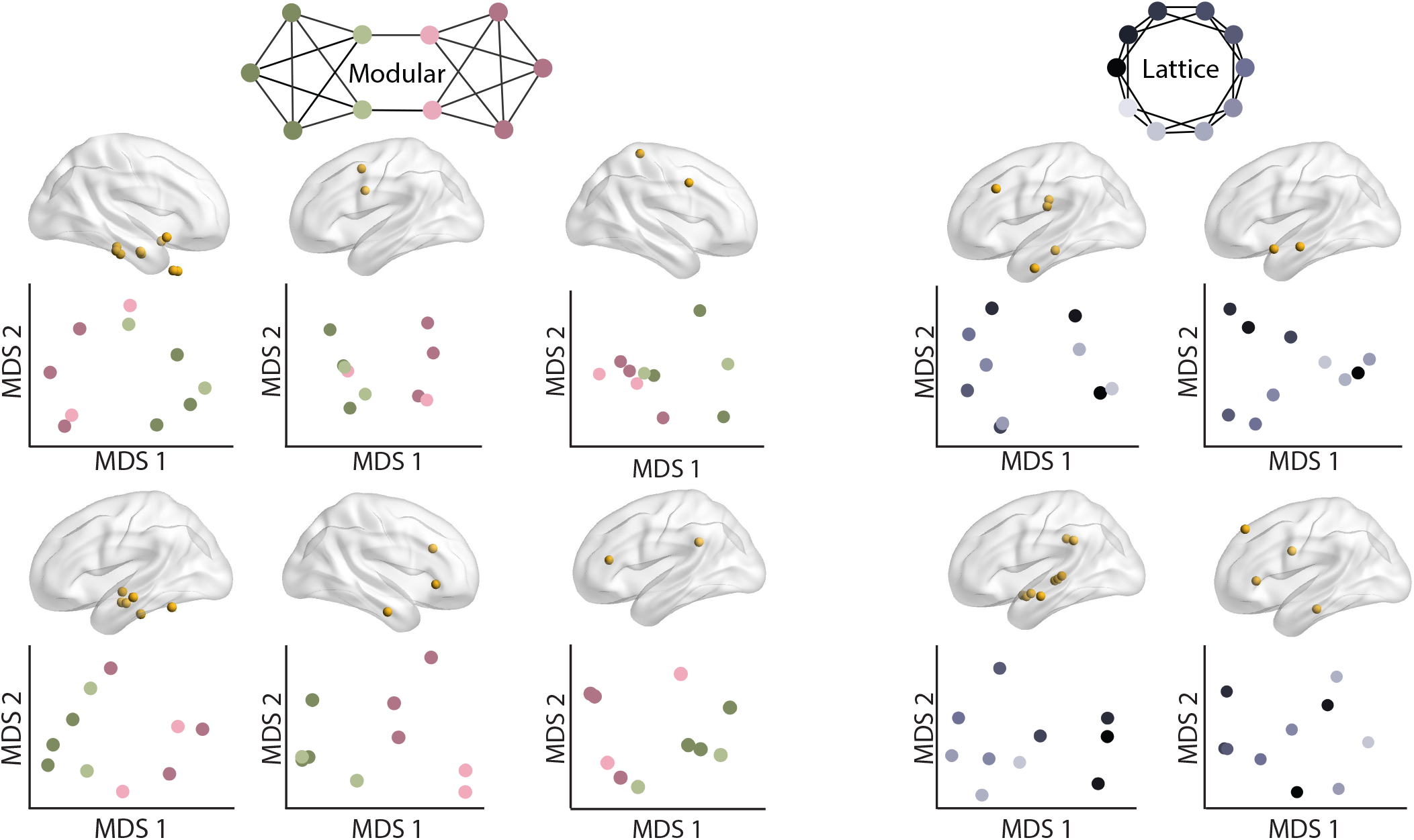
Diversity of low dimension projections of neural activity. The contacts used to create the underlying dissimilarity matrices (top). Low dimensional projections of neural dissimilarity matrices for each participant (bottom). For visualization purposes, only the hemisphere with the most contacts is shown. Participants who viewed walks from modular graphs are shown in the three leftmost columns, and participants who viewed walks from lattice graphs are shown in the two rightmost columns.

We next wished to test whether this modular separability in low dimensional neural activity is also present in latent spaces estimated from behavior. If modules are separable in both spaces, then temporally discounted space estimations might be sufficient to explain separability. If separability is only present in neural spaces, then further computations are likely needed to explain separability. We test for module discriminability using a linear discriminant analysis on low dimensional coordinates obtained from principal components analysis applied to neural distance matrices and on the estimated latent space. The linear discriminant analysis consists of training a linear classifier to label each node as being in the pink or green module based on the two dimensional coordinates. We then test how accurately that model predicts the true labels of the same data. The reported loss from this model is the proportion of stimuli that were misclassified. We find that for most participants, discriminability varied little between the data from neural dissimilarity and from the estimated latent space (**Fig. 5A**, paired *t*-test *t* = − 1.04, *p* = 0.344). However, 2 participants showed much lower discriminability for the estimated latent space, compared to the neural dissimilarity space (**Fig. 5A**). We next sought to test whether discriminability for estimated latent spaces was specific to a particular range of *β* values. We find that participants with higher *β* values show perfect discriminability whereas participants with lower *β* values do not (**Fig. 5B**). Visualization of the low dimensional projections from different *β* values shows that the poor discriminability was driven by the nodes with connections to other modules (**Fig. 5C**). More specifically, at a *β* value close to 0.1, we see an abrupt shift where nodes connecting two modules switch from being closer to their corresponding module, to being closer to the contrasting module. Taken together, these findings provide evidence for module discriminability in neural activity. However, whether that discriminability is predicted by the estimated latent space alone depends on the participant and diminishes for those participants characterized by low *β* values.

**FIG. 5.**
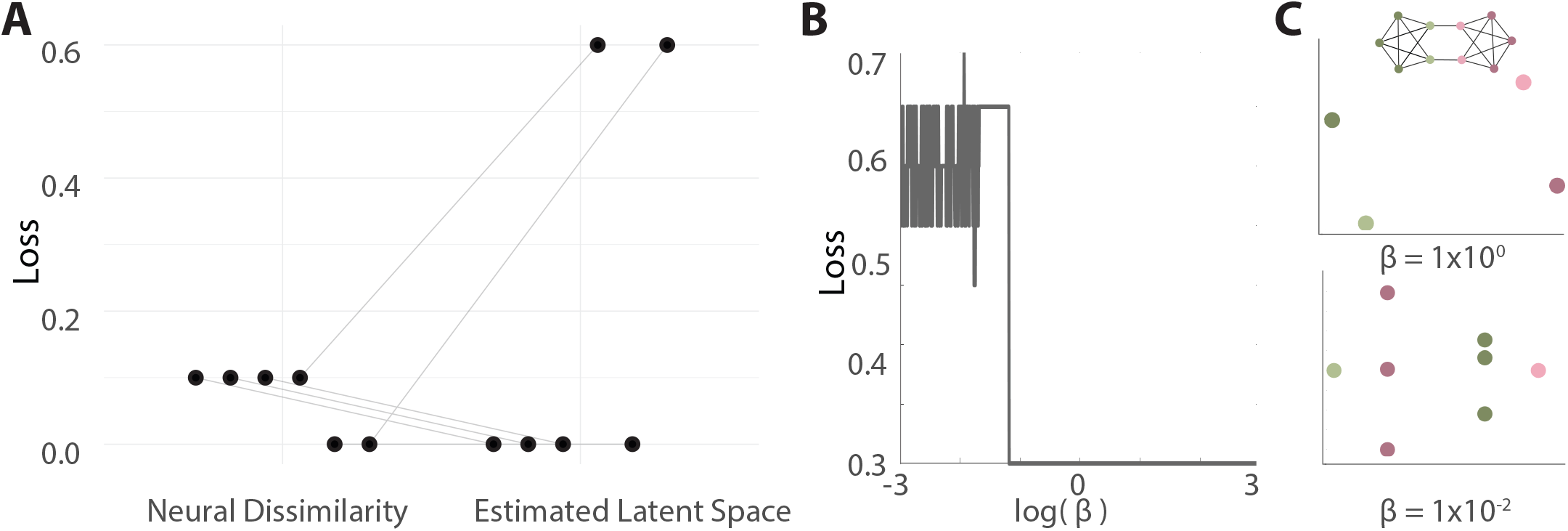
Module discriminability. *(A)* Loss, or incorrectly classified nodes, for each participant’s neural distance matrix and their estimated latent space. Lines connect the loss of the neural dissimilarity and estimated latent spaces for a single participant. *(B)* The loss for estimated latent spaces at different *β* values. *(C)* Visualization of a low dimensional projection of an estimated latent space from a large *β* value (top) and from a small *β* value (bottom) for one participant. Both plots show 10 points, though there is a high amount of overlap in the top plot, making only 4 points clearly distinguishable by eye.

### Temporal dynamics of latent space formation

In a final investigation, we sought to model how the estimated latent space might change during learning, and test whether neural activity showed similar temporal patterns. Assuming that *β* values are static during the course of learning, we can simulate how the estimated latent space *Â* changes on each trial due to each new transition observed between stimuli. We then calculated the correlation between the current estimated latent space at each trial *Â*(*t*) and the estimated latent space obtained using the infinite trial limit *Â* (**Fig. 6A**). Since participants only observed a finite walk, the quantity *Â*(*t*) does not converge to exactly *Â*. However, most participants quickly show high agreement between the finite and infinite-time estimates as they learn. Qualitatively, we see that larger *β* values result in a faster convergence towards the final *Â* regardless of the graph type (**Fig. 6A-B**).

**FIG. 6.**
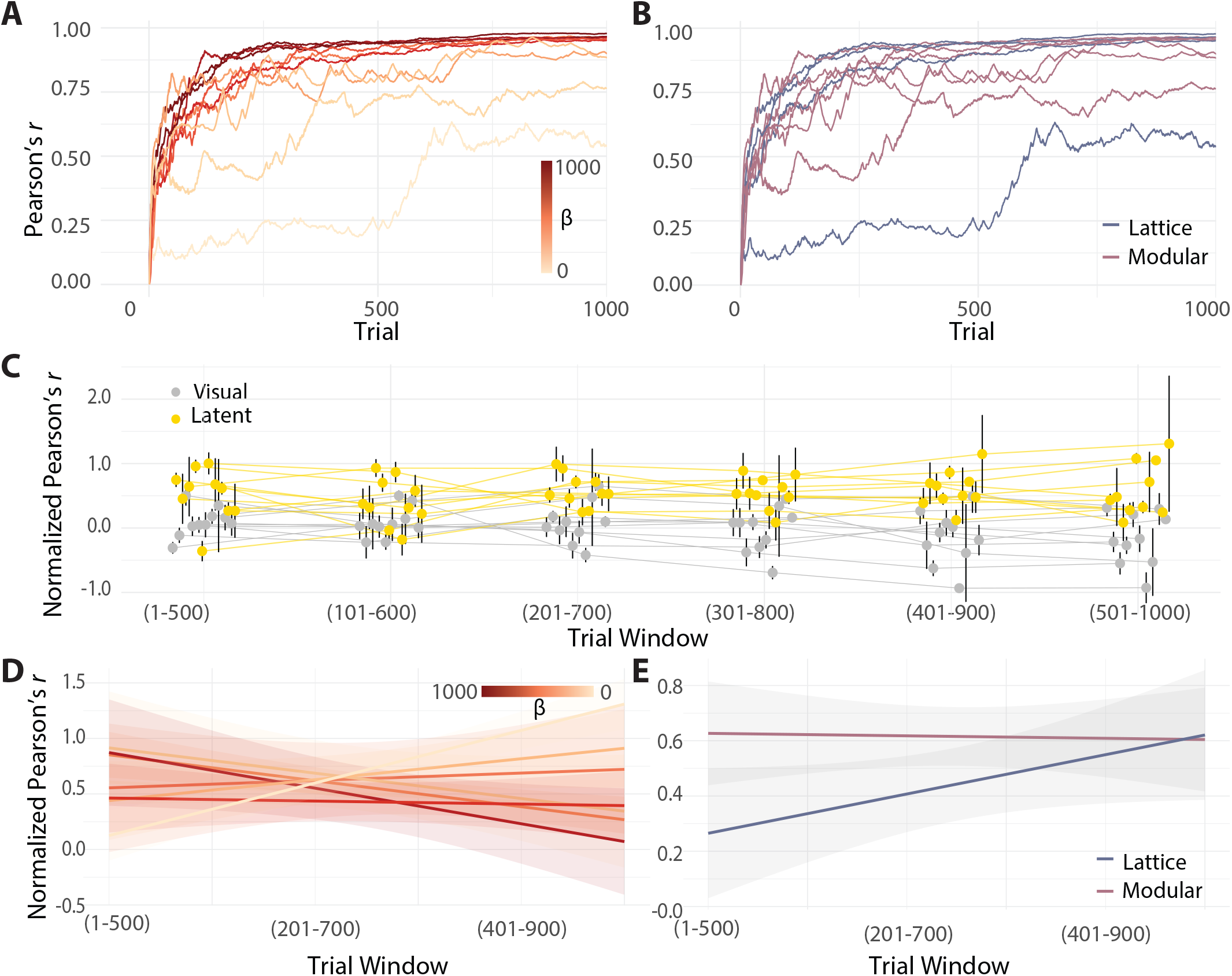
Time varying estimates of latent spaces. *(A)* The correlation between the infinite-time estimate of the latent space *Â* and the finite-time estimate of the latent space *Â*(*t*) at each trial *t*. Estimates with different *β* values are shown in different shades of orange. *(B)* The same information as that displayed in panel *(A)*, but now colored by graph type. Walks from lattice graphs are shown in blue and walks from modular graphs are shown in pink. *(C)* The correlation between the neural activity dissimilarity matrices and two spatial templates in two blocks of 500 trials. The correlation to the estimated latent space template is shown in yellow, and the correlation to the visual template is shown in grey. Error bars indicate the standard error of correlations over contacts. *(D)* Change in correlation of neural dissimilarity matrices to the latent space as a function of trial window, colored by the *β* value of the participant. Each line shows a linear fit of one participant’s change in correlation over time. Shaded regions indicate the 95% confidence interval. *(E)* The same information as that displayed in panel *(D)*, but now separated by graph type rather than by *β* value.

Informed by these data, we hypothesized that neural activity structure would also reflect the estimated latent space *Â* fairly early during learning. In order to ensure that we had enough trials to get stable estimates of activity structure, we tested this hypothesis by recalculating the correlation between the latent space and neural dissimilarity matrices in sliding blocks of 500 trials with a 100 trial offset. This process provided a total of 6 blocks. We recomputed these correlations only in individual contacts (*n* ranged from 2 to 10) whose activity was determined to be similar to the estimated latent space (see **Fig. 3**). We also calculated the correlation to the visual space in these same contacts as a comparison (**Fig. 6C**). Since we wished to capture the dynamics of contacts converging to their final values rather than differences in those final values, we normalized all correlation coefficients to the values calculated using all trials. We then averaged similarity values over all contacts and used a linear mixed-effects model to assess whether participants’ neural activity was more similar to the estimated latent space than the visual space, and if that similarity grew over time. In line with our hypothesis, we found significantly larger increases in correlation coefficients between the neural space and the latent space than in correlation coefficients between the neural space and visual space (linear mixed-effects model *F*_*window∗space*_ = 6.755, *p*_*window∗space*_ = 0.011), even in the first 500 trials (paired *t*-test *t* = 3.81, *p* = 0.004).

We next asked whether these changes in similarity were modulated by *β* values or by graph type. Our simulations suggest that participants with larger *β* values should show greater similarity to the latent space early during learning. Accordingly, we tested whether *β* values predicted the magnitude and rate of change of the normalized correlation coefficients between neural activity and the estimated latent space. We found significant changes in the magnitude of the normalized correlations associated with *β* values, as evidenced by a significant interaction between *β* values and the change in similarity over blocks (linear mixed-effects model *F*_*β*_ = 2.65, *p*_*β*_ − 0.110, *F*_*window∗β*_ = 6.12, *p*_*window∗β*_ = 0.017) (**Fig. 6D**). Specifically, smaller *β* values have positive relationships between the neural similarity to the estimated latent space and the trial window (increasing with learning), whereas larger *β* values have negative relationships. While the observed variation in the convergence of neural data to *Â* by *β* value was recapitulated in our simulations, we saw an additional trend towards less convergence over time that is not present in the model.

We also wished to determine whether changes in similarity (between neural activity and the latent space) over time were consistent across graph types, as predicted by our simulations. We found a significant interaction between the change in similarity over blocks and the graph type, which was not predicted by our model (linear mixed-effects model *F*_*graph*_ = 5.06, *p* = 0.029) (**Fig. 6E**). Participants who saw sequences from lattice graphs tended to have more positive slopes, whereas participants who saw sequences from modular graphs tended to have more negative slopes. It is worth noting that this separation by graph type reflects the significant interaction observed between reaction time and graph type in this same cohort.

Lastly, we sought to examine whether the discrepancies between our simulations and observations could be narrowed by relaxing the assumption that *β* values remain static during learning, and instead hypothesizing that finite-time estimates of *β* per block would diverge more from the infinite-time estimates of *β* in the later windows. To test this hypothesis, we recalculated *β* values for each participant in the same blocks of 500 trials. We then tested whether *β* values changed consistently across the population over blocks of trials. While we observed substantial variability in *β* values for some participants, there were no consistent differences across the population (**Fig. S6**, linear mixed-effects model *F*_*window*_ = 0.14, *p*_*window*_ = 0.714).

## DISCUSSION

In this work, we sought to better understand the neural correlates of latent space estimation from temporal sequences of stimuli that evince particular transition probability structures encoded as graphs. We utilized behavioral modeling to identify individual variations in temporal discounting and iEEG data recorded during learning to answer four main questions: (1) Do individuals in our iEEG cohort show behavioral evidence of learning an estimate of the latent space? (2) Which brain regions have neural activity that reflects these estimates? (3) Does the structure of neural activity facilitate the identification of task-relevant features? (4) Upon what time scale does neural structure appear, and is that timescale modulated by temporal discounting or graph structure? To answer question (1), we first had participants respond to cues generated from 2 different latent spaces: one with a modular structure, and one with a lattice structure. We found evidence that our iEEG cohort became faster and more accurate over time, consistent with participants learning the latent space and better anticipating upcoming stimuli. To answer question (2), we fit a model of learning that utilizes temporal discounting during latent space estimation, and found regions where neural activity has a similar structure to these estimates. For stimulus-evoked activity, most regions identified—regardless of the graph used to generate the sequences—were located in the temporal lobe, with some additional involvement of frontal structures.

Previous work investigating Euclidean spatial representations found that low dimensional projections of the estimated space readily identified task-relevant features like boundaries and modules[22]. This work motivated us to ask question (3), and accordingly to test whether there was evidence for the same identification of modules in our neural data. We found that for each participant who saw sequences drawn from a modular graph, low dimensional projections of neural activity in the selected temporal and frontal regions accurately separated each module, misidentifying at most one stimulus. Interestingly, this separability was not achieved as consistently in the estimated latent space itself, suggesting the possibility that neural processing enhances the separability of task-relevant features such as modules. Lastly, we leveraged the neural recordings taken during latent space learning to ask question (4), and accordingly to test predictions about how quickly participants acquire their estimates of the latent space. Our model predicted that estimates of the latent space would be formed within the first 500 trials, and that participants with stronger temporal discounting would converge faster. We found evidence in support of these hypotheses, and also additional differences in latent space learning based on graph type that were not predicted by our model. Ultimately, we determined where and when neural activity during a sequential reaction time task reflects individual variation in behavior, and how that activity related to recent theories that extend concrete cognitive maps to abstract spaces.

### Insights into probabilistic sequence learning

Previous work in probabilistic sequence learning has demonstrated that participants reacting to cues drawn from a random walk on a graph become sensitive to features of latent structure for a wide variety of graphs, with different numbers of stimuli, and across different sensory domains[5, 6, 10, 35, 54, 55]. Here, we significantly extend this literature by adapting a version of these tasks for use in patient populations with iEEG recordings. Using our adapted task, we find that both a healthy cohort recruited via Amazon’s Mechanical Turk and an iEEG cohort show evidence of learning, albeit with some differences in the nature of that learning.

We found that our mTurk cohort shows significant decreases in reaction time with increasing trial number, while our iEEG cohort shows decreases only across longer timescale blocks of 250 trials. While learning rates varied across the two cohorts, iEEG patients still performed the task with high accuracy (**Fig. 2A** and **B**), as expected given their cognitive capacities[53]. The slower learning is consistent with other work demonstrating poorer task performance in participants with epilepsy compared to controls[53, 56]. Patients with drug-resistant epilepsy were shown to have statistically significant decreases in task performance assessing motor function and cognitive attention[57], both of which are required for our experiment. However, our 2 cohorts are not matched on demographics or testing environment, making it difficult to determine whether differences are due to underlying epilepsy-related cognitive deficits or other factors.

We next tested for evidence of learning based on an increase in accuracy over time. While the mTurk cohort had relatively high accuracy throughout the experiment, their performance significantly decreased over time. While increasing speed and decreased accuracy are not necessarily indicative of disengagement from the task[58], this finding raises the possibility that some of the observed decreases in reaction time might be due to a decrease in correct responses. One possible explanation is a decrease in cognitive demand and arousal, leading to task disengagement and lower accuracy[59, 60]. In contrast to the mTurk cohort, the iEEG cohort shows an initial increase in accuracy, followed by a decrease in accuracy during later trials. Individuals with temporal lobe epilepsy tend to perform worse on tasks that demand higher order cognition and attention [61], and therefore it may be easier to engage with simpler tasks. While this quadratic relationship with accuracy still suggests a lower engagement with the task as time goes on, it corroborates the conclusion that the task is better suited for the iEEG cohort.

Lastly, we tested for differences in reaction time based on graph type. Previous research has found that participants tend to react faster to cues drawn from modular graphs than to those drawn from lattice or random graphs[6]. We do not recapitulate this finding in our mTurk population, possibly because the task we used was significantly simpler than that previously employed in Ref. [6], and hence did not entail the same learning complexity. There were also design differences between previous work and the current study; for example, here we used simpler motor commands (using one rather than two fingers at a time), fewer trials, breaks with rewarding feedback, and fewer unique stimuli. In our iEEG cohort, we found different rates of learning based on underlying graph type. However, future work with either a more complex task or a larger number of subjects is necessary to further validate this result.

### Insights into neural involvement in latent space estimation

To complement our study of behavior, we next probed the neural correlates of latent space estimation. We identified regions whose activity has a structure most similar to each individual’s estimated latent space (rather than to the true latent space). In performing this identification, we used a short window of activity locked to the stimulus, to the response, or to the middle of all trials. In contrast to the slower temporal resolution of metabolic neuroimaging techniques, iEEG allows for the use of short temporal windows to investigate neural activity structure in a time-resolved manner thereby providing insight not only into where, but also into when, structural representations emerge. We also compared 2 different similarity matrices to rule out possible alternative explanations of the observed structure that were unrelated to the estimated latent spaces. The first is a null model that takes the empirical trial data, and reorders it around a single point. Unlike shuffling trial order, this model preserves autocorrelative features of the data, and ensures that the observed similarity is specific to the observed walk sequence[51]; the second comparison is to a lower-level feature of stimulus appearance: the visual distance between highlighted stimuli on the screen. We expected this structure to be reflected in neural activity, and indeed many regions included contacts that were similar to both latent and visual spaces. Including these comparisons allows us to assess the selectivity of regional activity for structural, rather than visual, information.

#### Stimulus and response-locked activity implicate different brain regions

We found that for stimulus-locked activity, the most common regions identified were in the lateral, medial, and inferior temporal lobes. It is important to note that the temporal lobes also have more electrode coverage, and the identified regions made up between 4.6% and 27.3% percent of contacts in those areas; though not all highly sampled areas (e.g., superior temporal lobe) showed any contacts that were similar to either space. The presence of structure in this early evoked response is consistent with work demonstrating that changes in tuning curves of neurons in the medial temporal lobe (in both human and non-human primates) reflect statistical similarities between stimuli[62, 63]. For response-locked contacts, common areas still include the fusiform gyrus, but also include the inferior frontal gyrus, somatomotor area, and insula. This anatomical distribution is consistent with work showing that later stages of processing involve frontal regions receiving structural information from medial temporal regions. Further, the involvement of motor regions is certainly intuitive during response planning[64, 65].

#### Amygdala involvement in cognitive map formation

The region with both the highest percentage of contacts identified and the highest selectivity for the latent space was the amygdala, followed by the middle temporal lobe. The amygdala is a region often associated with processing of emotional and rewarding stimuli, and is highly connected to the hippocampus, with which it interacts during emotional memory[66]. Notably, some previous work using single unit human iEEG recordings has also shown activity reflective of cognitive map building in the amygdala[67, 68]. For example, in a study of single cell place selectivity in patients undergoing iEEG recording, the hippocampus demonstrated the most place-selective activity, yet cells in other parts of the medial temporal lobe, including the amygdala, showed selectivity as well[68]. Additionally, non-human primate studies have shown representations of abstract contexts for non-emotional stimuli in the amygdala[69]. Ultimately, our results corroborate these findings that amygdala activity can reflect abstract spaces.

#### Middle temporal lobe involvement in cognitive map formation

The second region identified, the middle temporal lobe, has also been identified in other iEEG studies of statistical learning. Previous work studying lower-level statistical learning using iEEG also identified primarily lateral temporal cortex, and little involvement of the hippocampus and entorhinal cortex[70]. Much work in human iEEG and fMRI implicates a broader range of temporal regions than comparable work in rodents[5, 16, 70]. This trend is likely partially due to the different cognitive and behavioral demands between species, but also raises the possibility of compensatory mechanisms in cohorts undergoing iEEG monitoring due to pathology in medial temporal lobes. This possibility cannot be ruled out completely, and therefore findings should ideally be corroborated in recordings from a healthy population. Some evidence of the role of lateral temporal lobe activity in learning a latent space from sequences exists in non-epileptic populations. Specifically, fMRI studies using similar tasks have also identified the interior temporal cortex to be reflective of some features of higher-order structure, but not reflective of the estimates of latent spaces as a whole[5].

#### Medial temporal lobe involvement in cognitive map formation

Much of the work in rodent and human latent space learning has focused on the hippocampal and entorhinal cortices, rather than lateral temporal lobes and amygdala[16]. Here, the hippocampus and sublobar temporal white matter are both implicated in our similarity analysis, supporting evidence of their important role in latent space learning. However, these areas are less common and less selective than lateral temporal structures and the amygdala in our data.

#### Other brain regions’ involvement in cognitive map formation

While the most common regions identified in our study and in previous work were in the temporal or frontal lobes, we observed multiple contacts in a wider distributed set of regions, including the insula, supplementary motor area, and precentral gyrus. Frontal areas, especially those in the medial prefrontal and orbitofrontal cortices, have been implicated in latent space learning, and are thought to be required at later stages than are medial temporal regions[71–73]. Consistent with these observations, we found that response-locked activity shows more involvement of these frontal areas. Additionally, some work in humans has shown that activity that reflects the estimated latent structure (e.g., in place and grid cells) is much more spatially distributed than in rodents, leading to theories that most of the cortex is actually capable of forming these representations[33, 74]. Our results are in line with these theories, and support the conclusion that diverse brain regions could support temporally discounted estimates of latent space. Taken together, neural activity most represented latent space estimates in the amygdala and middle temporal lobe when locked to the stimulus, whereas they most represented latent space estimates in the supplementary motor area and inferior frontal gyrus when locked to the response. These observations indicate that brain representations of learning are spatially distributed.

### Importance of low dimensional separation of task features

Studies investigating representations of spatial environments have pointed out the usefulness of low dimensional representations for learning to navigate[18, 22]. Evidence for dimensionality reduction of neural signals has been observed in neural structures at 3 distinct scales: single neurons, anatomical regions, and the whole brain[75–78]. Broadly, dimensionality reduction of neural signals is thought to enable the brain to easily extract important, often changing information and facilitating the development of a sparse, efficient neural code for items in the environment[77, 79]. For dimensionality reduction of cognitive maps specifically, much work has focused on the medial temporal lobe. For example, the hippocampus has been functionally modeled as a variational autoen-coder that continuously compresses incoming structural and sensory information to identify similar contexts[18]. Additionally, properties of grid cells[22], commonly but not exclusively found in the entorhinal cortex[80], can be explained by the eigenvectors of a temporally discounted estimate of the latent space. Importantly, these low dimensional bases identify borders and modules in simulated spaces, the same features thought to be useful for successful navigation[22].

We asked whether these modeling observations were recreated in an abstract relational space. Using linear discriminant analysis, we found that modules are highly discriminable in individuals who saw sequences drawn from a modular graph. Interestingly, many of the estimated latent spaces show the same level of discriminability, although some show levels far lower. Upon further investigation, we find that the discriminability of estimated latent spaces was determined by the associated *β* value. We chose linear discrimination as a conservative estimate of separability, which is biologically implementable in theory by few neurons whose firing mimics the low dimensional bases[81]; however, other methods of identifying modules are theoretically possible[82].

The discussion of these findings raises the possibility that neural systems are transforming or building estimates of latent spaces in a way that enhances the separability of modules. One hypothesis is that the increased separability in low dimensional space arises from neurons with high dimensional, combinatorial responses to individual stimuli[82]. These types of neurons are thought to be present in associative areas such as the frontal cortex and medial temporal lobes[82]. It is hence intuitively plausible that regions in lower-order areas are less able to separate modules, but potentially more able to distinguish individual stimuli[82]. While these theories are based on the function of individual neurons, similar ideas can be extended to neuronal populations. Accordingly, future work could test whether divergences of neural activity from the estimated latent space increase the separability of modules at all places on the neural processing hierarchy, or only at more transmodal areas. The observation that neural dissimilarity better separates modules than the corresponding latent space estimation presents interesting directions for further investigation independent of validations of those theories.

### Limitations

Here, we have put forth new evidence for neural correlates of latent space learning, although these results should be interpreted in light of the various limitations of our study. Many of our analyses focused on individual participants, an approach that is especially well-suited for iEEG analysis given the small and heterogeneous samples. However, some results—including evidence for learning and temporal changes in neural similarity structure—were assessed at the group level. To supplement these findings, we also present larger behavioral cohorts and numerical simulations. Despite these techniques, our group level results would be further strengthened by replication in larger samples.

Additionally, we sought to identify the regions whose activity was structured most similarly to the estimated latent space. This involved selecting contacts with stronger correlations than 95% of null models. This selection process means that there is a chance that some contacts would be retained due to basic features of neural activity, and not due to task structure. Because of this fact, we highlight the regions where multiple contacts were identified, reducing the likelihood that our conclusions depend on false positives. We approached identifying contacts with activity structure similar to the latent space in a data driven manner, and therefore expected the same pattern of activity in all regions. We also grouped all identified regions together when investigating properties of low dimensional projections of neural activity structure. However, there is good evidence that specific regions, or even locations within the same region might be active at different times[72] or use slightly altered transformations of the estimated space[29]. Identifying these differences is an important pathway for future analysis, but would require a larger cohort, where more individuals reliably show activity in the regions of interest, or a hypothesis-driven rather than data-driven assessment of regional contributions.

Lastly, we show evidence that some predictions of how latent spaces are learned over time are borne out in neural data. However, our model only uses one relatively simple learning rule. Other work has tested a variety of learning rules that all give rise to temporally discounted latent space estimations, and has shown that some are more consistent with neural activity than others[83]. Here, we do not intend to claim that the implemented rule was more accurately reflecting changes in neural activity than others, but simply to identify the ways that estimated latent spaces appear in neural activity. Future work investigating and comparing different learning rules would be a welcome contribution to the field.

### Future directions

Studying latent space learning presents an exciting opportunity in neuroscience to connect theoretical models to both behavior and hypothesized neural mechanisms for the implementation of these models. Work in rodents has suggest that temporally discounted estimates of relational spaces are built through synchronization of cell populations to *θ* rhythms (4-10 Hz)[84]. Distinct populations of cells in the CA1 subfield of the hippocampus synchronize their firing to the peaks of *θ* rhythms; the firing of different cells then becomes bound together via plasticity to represent unique temporal contexts[85]. Within the hippocampus, map-like firing patterns of these linked assemblies of neurons reflect physical relationships after exploring new environments[86]. The phase of *θ* rhythms also synchronizes with activity in cortical areas such as the prefrontal cortex where information about temporal context is used for other processes[65]. In humans, *θ* rhythms have been implicated in tasks requiring estimates of an underlying latent space, including episodic memory, spatial navigation, and semantic memory[87]. However, there is also evidence that these rhythms are less important for human learning than for rodent learning, and some investigators even hypothesize that other mechanisms, such as saccades, are responsible for the synchronization of cell populations[16]. Similar studies to clarify the role of *θ* rhythms during latent space learning would extend the field appreciably.

Beyond connections to mechanistic neural implementations of these models, further extensions to more ecological contexts would also benefit our understanding of latent space learning, and how they influence diverse cognitive processes. Extensions of this theory to ecological network structures, and to different exploration strategies and walk types have already been discussed and implemented[35, 88, 89]. Nevertheless, our work suggests that further advancements could expand the theory to incorporate temporal variability in learning strategies. Here, we show preliminary evidence for a change in learning rates based on the extent of temporal discounting, and also a shift in the extent of temporal discounting used over time (**Fig. S6**). One would expect that different amounts of temporal discounting might be better suited to different tasks, tasks occurring at different timescales, or even different stages of the same task. This intuition is consistent with work demonstrating that different brain regions, or even different parts of the hippocampus, are sensitive to different timescales of information[72], which could potentially be related to different amounts of temporal discounting. Extension of this work to incorporate more dynamic models of learning would help us better understand domain-general latent space learning, and further align the models of these behaviors with the evidence of their implementation in the brain.

## CITATION DIVERSITY STATEMENT

Recent work in several fields of science has identified a bias in citation practices such that papers from women and other minority scholars are under-cited relative to the number of such papers in the field [90**? ? ? ?** –94]. Here we sought to proactively consider choosing references that reflect the diversity of the field in thought, form of contribution, gender, race, ethnicity, and other factors. First, we obtained the predicted gender of the first and last author of each reference by using databases that store the probability of a first name being carried by a woman [94, 95]. By this measure (and excluding self-citations to the first and last authors of our current paper), our references contain 10.55% woman(first)/woman(last), 15.43% man/woman, 18.83% woman/man, and 55.18% man/man. This method is limited in that a) names, pronouns, and social media profiles used to construct the databases may not, in every case, be indicative of gender identity and b) it cannot account for intersex, non-binary, or transgender people. Second, we obtained predicted racial/ethnic category of the first and last author of each reference by databases that store the probability of a first and last name being carried by an author of color [96, 97]. By this measure (and excluding self-citations), our references contain 9.17% author of color (first)/author of color(last), 10.93% white author/author of color, 20.72% author of color/white author, and 59.18% white author/white author. This method is limited in that a) names and Florida Voter Data to make the predictions may not be indicative of racial/ethnic identity, and b) it cannot account for Indigenous and mixed-race authors, or those who may face differential biases due to the ambiguous racialization or ethnicization of their names. We look forward to future work that could help us to better understand how to support equitable practices in science.

## I. AUTHOR CONTRIBUTIONS

J.S., V.N.R., and D.S.B planned analyses. J.S. and C.W.L. wrote code. J.S., A.E.K., and D.S.B. designed the behavioral experiment. J.S. and A.E.K. collected behavioral data from Amazon Mechanical Turk. A.R. and J.M.S. localized contacts to atlases. J.S., K.P.S, and R.A. collected iEEG data. All authors contributed to writing the manuscript.

## ACKNOWLEDGEMENTS

We would like to thank Dale Zhou for helpful discussions regarding dimensionality reduction in the brain. We also acknowledge the help of Everett Prince, Jacqueline Boccanfusco, Magda Wernovsky, Amanda Samuel, and all the employees at the CNT who make the collection and curation of iEEG data possible. D.S.B. and J.S. acknowledge support from the John D. and Catherine T. MacArthur Foundation, the Alfred P. Sloan Foundation, the ISI Foundation, the Paul Allen Foundation, the Army Research Office (Grafton-W911NF-16-1-0474, DCIST-W911NF-17-2-0181), the National Institute of Mental Health (2-R01-DC-009209-11, R01 – MH112847, R01-MH107235, R21-M MH-106799), National Institute of Neurological Disorders and Stroke (R01 NS099348), and the National Science Foundation (BCS-1631550). J.S. acknowledges support from the National Institute of Mental Health National Research Service Award (F31MH120925). The content is solely the responsibility of the authors and does not necessarily represent the official views of any of the funding agencies. Lastly, we would like to thank the patients and their families for their participation and support.

