## supplement for "Neurophysiological evidence for cognitive map formation during sequence learning"

[1] *Department of Bioengineering, School of Engineering and Applied Science, University of Pennsylvania, Philadelphia, PA 19104, USA*
[2] *Initiative for the Theoretical Sciences, Graduate Center, City University of New York, New York, NY 10016, USA*
[3] *Joseph Henry Laboratories of Physics, Princeton University, Princeton, NJ 08544, USA*
[4] *Princeton Neuroscience Institute, Princeton University, Princeton, NJ 08544, USA*
[5] *Department of Neurology, Hospital of the University of Pennsylvania, Philadelphia, PA 19104, USA*
[6] *Department of Radiology, Hospital of the University of Pennsylvania, Philadelphia, PA 19104, USA*
[7] *Department of Electrical & Systems Engineering, School of Engineering & Applied Science, University of Pennsylvania, Philadelphia, PA 19104, USA*
[8] *Department of Neurology, Perelman School of Medicine, University of Pennsylvania, Philadelphia, PA 19104, USA*
[9] *Department of Psychiatry, Perelman School of Medicine, University of Pennsylvania, Philadelphia, PA 19104, USA*
[10] *Department of Physics & Astronomy, College of Arts & Sciences, University of Pennsylvania, Philadelphia, PA 19104, USA*
[11] *The Santa Fe Institute, Santa Fe, NM 87501, USA and*

| Participant | Sex | Age (yrs) | Race | Ephys | Median RT | Graph | Beta | Coverage |
|---|---|---|---|---|---|---|---|---|
| 1 | F | 36 | White | Y | 0.923 | Lattice | 0.024 | Left: medial, ventral, and lateral temporal; central gyrus/sulcus; insula; middle and inferior frontal. Right: medial, ventral, and lateral temporal. |
| 2 | M | 24 | White | Y | 0.546 | Modular | 0.17 | Right: medial, ventral, and lateral temporal; insula; middle and inferior frontal. |
| 3 | F | 25 | White | Y | 1.103 | Lattice | 1000 | Left: medial, ventral, and lateral temporal; inferior parietal; middle and inferior occipital; insula; cingulate. Right: medial, ventral, and lateral temporal; inferior parietal; central gyrus/sulcus; insula. |
| 4 | F | 32 | White | Y | 0.572 | Modular | 0.43 | Left: medial, ventral, and lateral temporal; inferior occipital; insula; inferior frontal. Right: medial and lateral temporal; insula. |
| 5 | F | 47 | White | Y | 0.657 | Lattice | 1000 | Left: medial, ventral, and lateral temporal; inferior parietal; middle and inferior occipital. |
| 6 | F | 58 | White | Y | 0.732 | Modular | 0.12 | Right: superior and middle frontal; central gyrus/sulcus; supplemental motor; cingulate. |
| 7 | M | 21 | White | Y | 0.724 | Lattice | 0.33 | Left: medial, ventral, and lateral temporal; cingulate; central and middle occipital; superior parieta; superior, middle, and inferior frontal. |
| 8 | M | 22 | White | Y | 0.482 | Modular | 0.024 | Left: medial, ventral, and lateral temporal; insula; middle and inferior frontal. |
| 9 | F | 22 | White | Y | 1.075 | Modular | 0.23 | Left: medial and ventral temporal; insula; inferior and middle frontal; basal ganglia. Right: medial, ventral, and middle temporal; inferior and superior frontal. |
| 10 | F | 37 | White | Y | 0.795 | Modular | 0.044 | Left: medial, ventral, and lateral temporal; insula; inferior frontal; cingulate. |
| 11 | F | 47 | Black | Y | 0.696 | Modular | 0 | - |
| 12 | F | 23 | White | N | 1 | Modular | - | - |
| 13 | F | 39 | White | N | 0.773 | Modular | - | - |

TABLE I. **Participant Demographics.** Demographic and task relevant information about each participant. The 'Ephys' column indicates whether or not the participant had electrophysiological recordings. The 'Median RT' column provides the median reaction time.

[1] A. E. Kahn, E. A. Karuza, J. M. Vettel, and D. S. Bassett, Network constraints on learnability of probabilistic motor sequences, Nature Human Behaviour 10.1038/s41562-018-0463-8 (2017), arXiv:1709.03000.

| AAL Region | Visual Count | Latent Count | Visual (%) | Latent (%) | Total Count |
|---|---|---|---|---|---|
| Amygdala | 1 | 3 | 9.09 | 27.2727273 | 11 |
| Angular | 1 | 0 | 11.11 | 0 | 9 |
| Calcarine | 2 | 0 | 28.57 | 0 | 7 |
| Caudate | 0 | 0 | 0 | 0 | 6 |
| Cerebellum_4_5 | 1 | 0 | 100 | 0 | 1 |
| Cingulum_Ant | 0 | 0 | 0 | 0 | 3 |
| Cingulum_Mid | 0 | 0 | 0 | 0 | 1 |
| Cingulum_Post | 0 | 1 | 0 | 16.6666667 | 6 |
| Cuneus | 0 | 0 | 0 | 0 | 1 |
| Frontal_Inf_Oper | 0 | 1 | 0 | 14.2857143 | 7 |
| Frontal_Inf_Orb | 0 | 1 | 0 | 8.33333333 | 12 |
| Frontal_Inf_Tri | 10 | 2 | 19.23 | 3.84615385 | 52 |
| Frontal_Med_Orb | 0 | 0 | 0 | 0 | 2 |
| Frontal_Mid | 3 | 1 | 16.61 | 5.55555556 | 18 |
| Frontal_Mid_Orb | 0 | 0 | 0 | 0 | 3 |
| Frontal_Sup | 1 | 1 | 10 | 10 | 10 |
| Frontal_Sup_Orb | 0 | 0 | 0 | 0 | 2 |
| Fusiform | 4 | 5 | 8.69565217 | 10.8695652 | 46 |
| Heschl | 0 | 0 | 0 | 0 | 2 |
| Hippocampus | 4 | 2 | 7.84313725 | 3.92156863 | 51 |
| Insula | 1 | 2 | 4.76190476 | 9.52380952 | 21 |
| Lingual | 0 | 0 | 0 | 0 | 5 |
| NotInAtl | 25 | 11 | 9.65250965 | 4.24710425 | 259 |
| Occipital_Inf | 1 | 0 | 20 | 0 | 5 |
| Occipital_Mid | 0 | 0 | 0 | 0 | 9 |
| Olfactory | 0 | 0 | 0 | 0 | 1 |
| Pallidum | 0 | 0 | 0 | 0 | 1 |
| ParaHippocampal | 1 | 1 | 4.34782609 | 4.34782609 | 23 |
| Parietal_Inf | 1 | 0 | 16.6666667 | 0 | 6 |
| Parietal_Sup | 0 | 0 | 0 | 0 | 1 |
| Postcentral | 0 | 1 | 0 | 9.09090909 | 11 |
| Precentral | 4 | 2 | 23.5294118 | 11.7647059 | 17 |
| Precuneus | 1 | 0 | 33.3333333 | 0 | 3 |
| Putamen | 0 | 0 | 0 | 0 | 5 |
| Rectus | 1 | 0 | 100 | 0 | 1 |
| Rolandic_Oper | 0 | 0 | 0 | 0 | 12 |
| Supp_Motor_Area | 0 | 0 | 0 | 0 | 5 |
| SupraMarginal | 0 | 0 | 0 | 0 | 7 |
| Temporal_Inf | 5 | 5 | 4.62962963 | 4.62962963 | 108 |
| Temporal_Mid | 4 | 6 | 3.30578512 | 4.95867769 | 121 |
| Temporal_Pole_Mid | 2 | 0 | 18.1818182 | 0 | 11 |
| Temporal_Pole_Sup | 1 | 0 | 20 | 0 | 5 |
| Temporal_Sup | 2 | 1 | 6.06060606 | 3.03030303 | 33 |

TABLE II. **Grey Matter Contacts.** The locations of all contacts in the AAL atlas.

| Tal. Region | Visual # | Latent # | Visual (%) | Latent (%) | Total Count |
|---|---|---|---|---|---|
| | 3 | 3 | 17.6470588 | 2.47933884 | 25 |
| Cerebellum.AnteriorLobe.Culmen.GrayMatter | 1 | 0 | 3.03030303 | 0 | 8 |
| Cerebellum.PosteriorLobe.Declive.GrayMatter | 1 | 0 | 9.09090909 | 0 | 6 |
| FrontalLobe.InferiorFrontalGyrus.GrayMatter.Brodmannarea45 | 1 | 0 | 9.09090909 | 0 | 5 |
| FrontalLobe.InferiorFrontalGyrus.WhiteMatter | 6 | 1 | 11.5384615 | 1.92307692 | 28 |
| FrontalLobe.MiddleFrontalGyrus | 2 | 0 | 1.65289256 | 0 | 3 |
| FrontalLobe.MiddleFrontalGyrus.GrayMatter.Brodmannarea6 | 0 | 1 | 0 | 0 | 3 |
| FrontalLobe.MiddleFrontalGyrus.GrayMatter.Brodmannarea9 | 1 | 0 | 11.1111111 | 0 | 3 |
| FrontalLobe.MiddleFrontalGyrus.WhiteMatter | 1 | 2 | 100 | 20 | 24 |
| FrontalLobe.ParacentralLobule.WhiteMatter | 0 | 1 | 0 | 0 | 2 |
| FrontalLobe.PrecentralGyrus.GrayMatter.Brodmannarea4 | 1 | 0 | 10 | 0 | 8 |
| FrontalLobe.PrecentralGyrus.GrayMatter.Brodmannarea6 | 2 | 2 | 11.1111111 | 11.1111111 | 9 |
| FrontalLobe.PrecentralGyrus.WhiteMatter | 1 | 0 | 4.76190476 | 0 | 16 |
| FrontalLobe.Sub-Gyral.WhiteMatter | 10 | 3 | 3.86100386 | 1.15830116 | 71 |
| FrontalLobe.Sub-Gyral.WhiteMatter.CorpusCallosum | 1 | 0 | 4.76190476 | 0 | 4 |
| FrontalLobe.SubcallosalGyrus.WhiteMatter | 1 | 1 | 20 | 4.34782609 | 3 |
| FrontalLobe.SuperiorFrontalGyrus.WhiteMatter | 0 | 1 | 0 | 0 | 3 |
| LimbicLobe.AnteriorCingulate.WhiteMatter | 1 | 0 | 4.34782609 | 0 | 8 |
| LimbicLobe.CingulateGyrus.WhiteMatter | 2 | 0 | 28.5714286 | 0 | 19 |
| LimbicLobe.ParahippocampalGyrus.GrayMatter.Amygdala | 1 | 2 | 16.6666667 | 66.6666667 | 10 |
| LimbicLobe.ParahippocampalGyrus.GrayMatter.Brodmannarea34 | 1 | 0 | 100 | 0 | 2 |
| LimbicLobe.ParahippocampalGyrus.GrayMatter.Hippocampus | 0 | 1 | 0 | 0 | 7 |
| LimbicLobe.ParahippocampalGyrus.WhiteMatter | 6 | 4 | 5.55555556 | 3.7037037 | 45 |
| LimbicLobe.PosteriorCingulate.GrayMatter.Brodmannarea23 | 0 | 1 | 0 | 0 | 1 |
| OccipitalLobe.FusiformGyrus.GrayMatter.Brodmannarea19 | 1 | 0 | 20 | 0 | 3 |
| OccipitalLobe.LingualGyrus.GrayMatter.Brodmannarea18 | 1 | 0 | 16.6666667 | 0 | 1 |
| OccipitalLobe.LingualGyrus.WhiteMatter | 1 | 0 | 14.2857143 | 0 | 6 |
| OccipitalLobe.MiddleOccipitalGyrus.GrayMatter.Brodmannarea19 | 1 | 0 | 8.33333333 | 0 | 1 |
| OccipitalLobe.MiddleOccipitalGyrus.WhiteMatter | 2 | 0 | 18.1818182 | 0 | 12 |
| ParietalLobe.SupramarginalGyrus.GrayMatter.Brodmannarea40 | 1 | 0 | 9.09090909 | 0 | 1 |
| ParietalLobe.SupramarginalGyrus.WhiteMatter | 1 | 1 | 0 | 0 | 12 |
| Sub-lobar.Claustrum.GrayMatter | 1 | 1 | 0 | 0 | 6 |
| Sub-lobar.Extra-Nuclear.GrayMatter.Brodmannarea13 | 0 | 1 | 0 | 0 | 1 |
| Sub-lobar.Extra-Nuclear.WhiteMatter | 4 | 3 | 8.69565217 | 6.52173913 | 53 |
| Sub-lobar.Extra-Nuclear.WhiteMatter.CorpusCallosum | 1 | 0 | 0 | 0 | 4 |
| Sub-lobar.Insula.WhiteMatter | 0 | 1 | 0 | 0 | 11 |
| Sub-lobar.LateralVentricle.Cerebro-SpinalFluid | 2 | 0 | 6.06060606 | 0 | 18 |
| TemporalLobe.FusiformGyrus.GrayMatter.Brodmannarea20 | 1 | 1 | 0 | 0 | 12 |
| TemporalLobe.FusiformGyrus.WhiteMatter | 1 | 3 | 0 | 0 | 19 |
| TemporalLobe.InferiorTemporalGyrus | 1 | 0 | 0 | 0 | 5 |
| TemporalLobe.InferiorTemporalGyrus.GrayMatter.Brodmannarea20 | 1 | 2 | 0 | 0 | 15 |
| TemporalLobe.InferiorTemporalGyrus.WhiteMatter | 1 | 0 | 0 | 0 | 22 |
| TemporalLobe.MiddleTemporalGyrus | 0 | 1 | 0 | 0 | 10 |
| TemporalLobe.MiddleTemporalGyrus.GrayMatter.Brodmannarea21 | 1 | 1 | 0 | 0 | 25 |
| TemporalLobe.MiddleTemporalGyrus.WhiteMatter | 1 | 4 | 0 | 0 | 67 |
| TemporalLobe.Sub-Gyral.GrayMatter.Brodmannarea21 | 0 | 1 | 0 | 0 | 1 |
| TemporalLobe.Sub-Gyral.GrayMatter.Brodmannarea37 | 1 | 0 | 0 | 0 | 2 |
| TemporalLobe.Sub-Gyral.WhiteMatter | 4 | 3 | 7.84313725 | 5.88235294 | 93 |
| TemporalLobe.SuperiorTemporalGyrus.WhiteMatter | 4 | 0 | 23.5294118 | 0 | 41 |
| TemporalLobe.SupramarginalGyrus.WhiteMatter | 1 | 0 | 0 | 0 | 2 |

TABLE III. **White Matter Contacts.** The locations of all contacts outside the AAL atlas were labeled by the Talairach atlas. Most are in deeper white matter structures. For display purposes, the ‘Cerebrum’ portion of the label has been removed from all cortical areas.

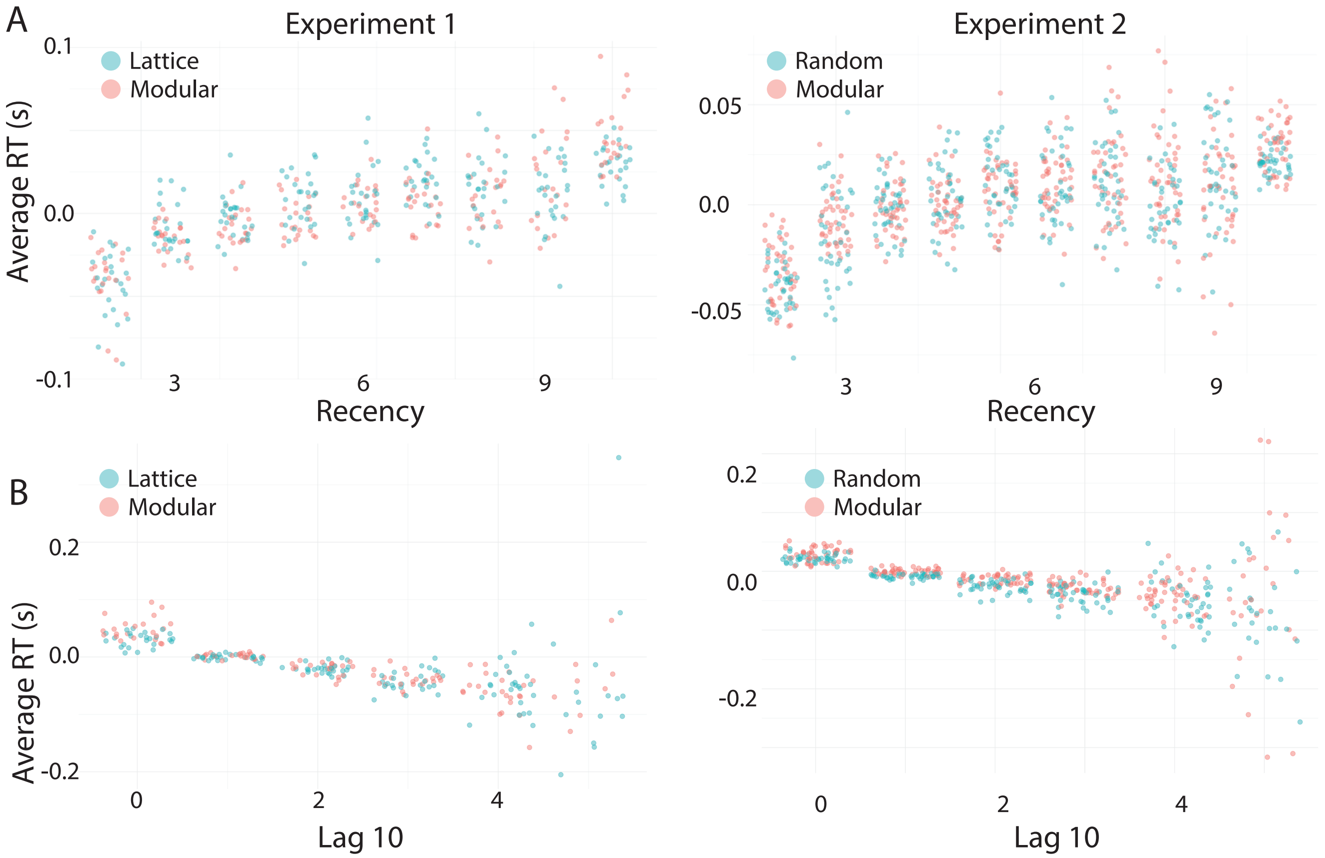


FIG. 1. **Comparison of Variables for Quantifying Recency.** *(A)* The average residual reaction times for each subject in each recency bin. The first bin (2) is the average of all reactions when the current stimulus was seen two stimuli ago. This data is for participants in the mTurk cohort of our paper, here labeled 'Experiment 1'. *(B)* The same as in panel *(A)*, but for the participants from Ref. [1], here called 'Experiment 2'. *(C)* The same as in panel *(A)*, but for lag10 bins. Here, the first bin (0) is the average reaction time to all trials where the current stimulus was seen 0 times in the past 10 trials. *(D)* The same as in panel *(C)*, but for the participants from Ref. [1].

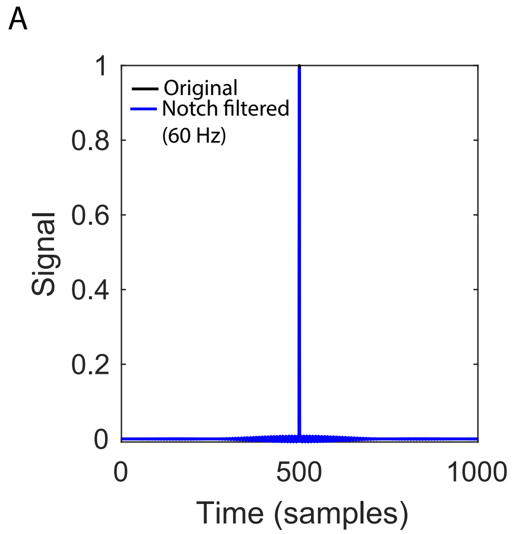


FIG. 2. **Notch Filter Response.** The response of the Notch filter used in the paper to a sharp transient signal. The original signal is shown in black, and the Notch filtered signal in blue. The signals are highly overlapping, indicating that perturbations to the signal resulting from filtering sharp transients are small.

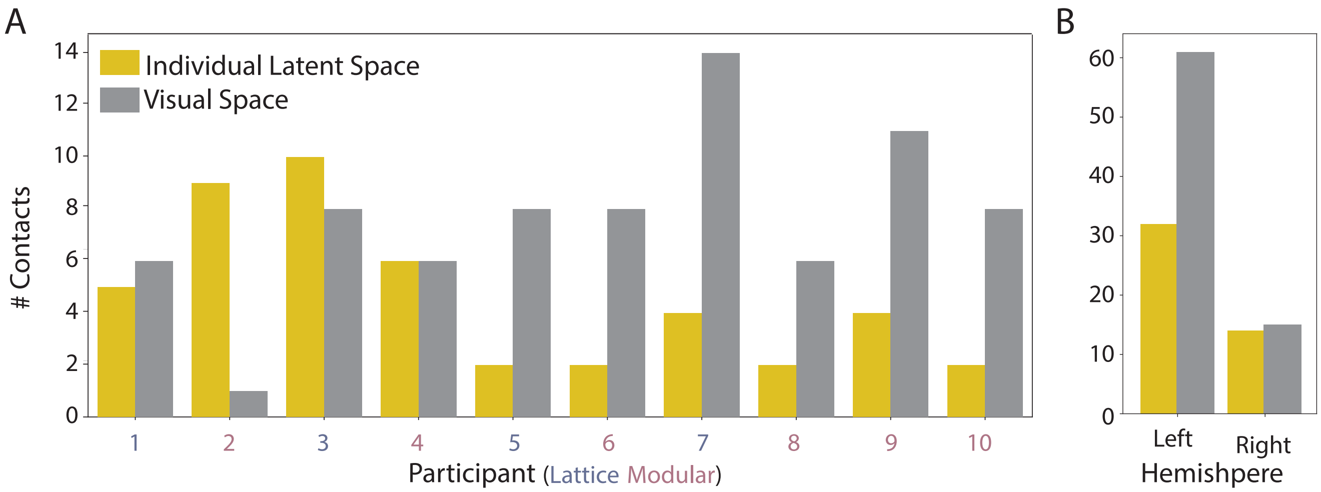


FIG. 3. **Additional Information About Stimulus-Aligned Contacts.** *(A)* The number of contacts retained for each individual for both the latent (gold) and visual (grey) space. *(B)* The number of contacts retained in each hemisphere for each space.

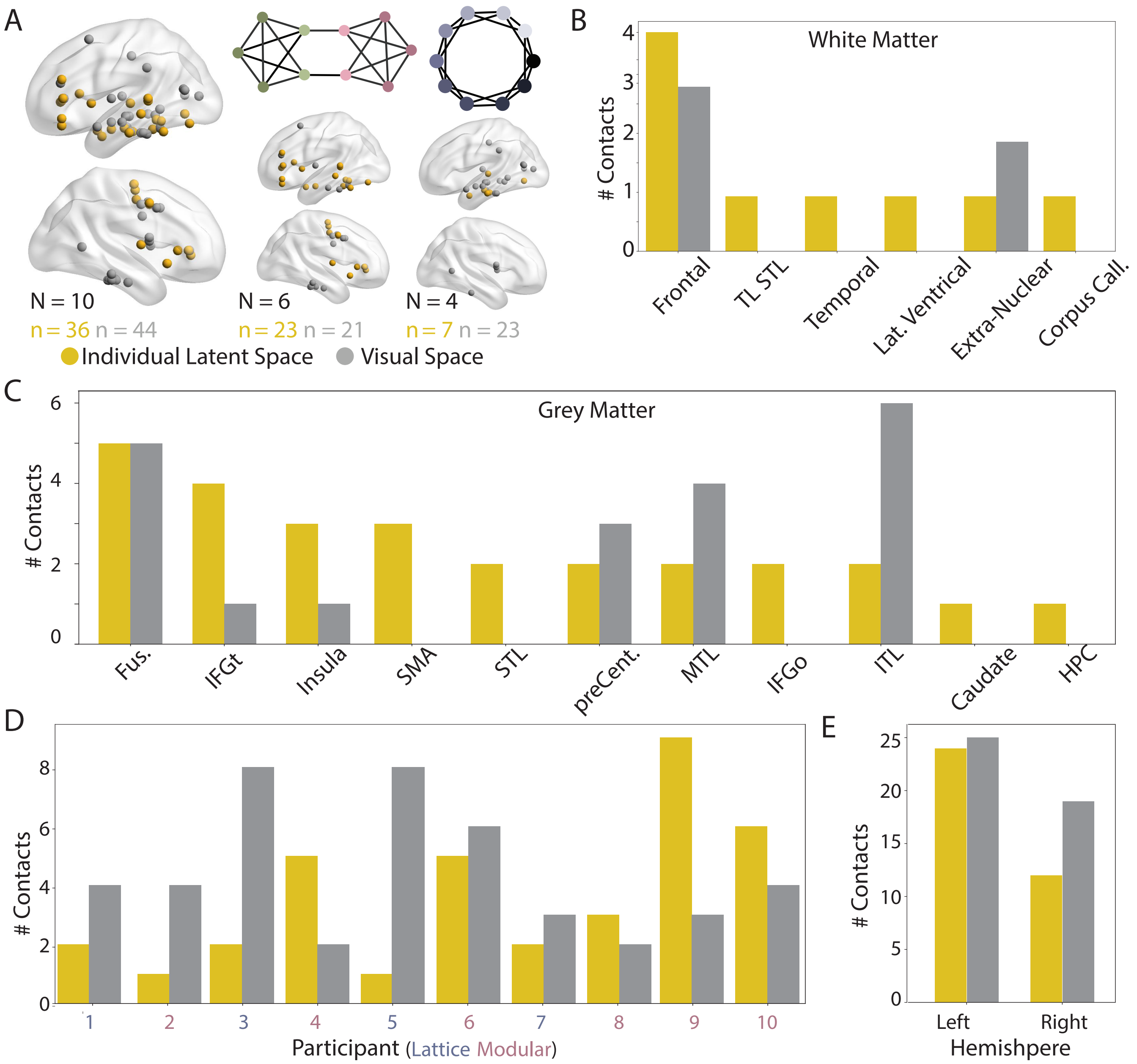


FIG. 4. **Response-Aligned Contacts.** *(A)* Visualization of retained contacts from response-aligned data with similar activity spaces on an MNI brain. Contacts that are similar to the latent space are shown in gold, and contacts that are similar to the visual space are shown in grey. *(B-C)* Anatomical localization of grey (*(B)*) and white (*(C)*) matter contacts. Grey matter contacts are localized with the AAL atlas, and white matter contacts are localized with the Talaraich atlas. *(D)* The number of contacts retained in each hemisphere for each space. *(E)* The number of contacts retained for each individual for each space. ITL: inferior temporal lobe, MTL: middle temporal lobe, STL: superior temporal lobe, TL STL: temporal lobe superior temporal lobe, Fus: fusiform, HPC: hippocampus, IFGt: inferior frontal gyrus (pars triangularis), IFGo: inferior frontal gyrus (pars orbitalis), SMA: supplemental motor area, preCent: precentral.

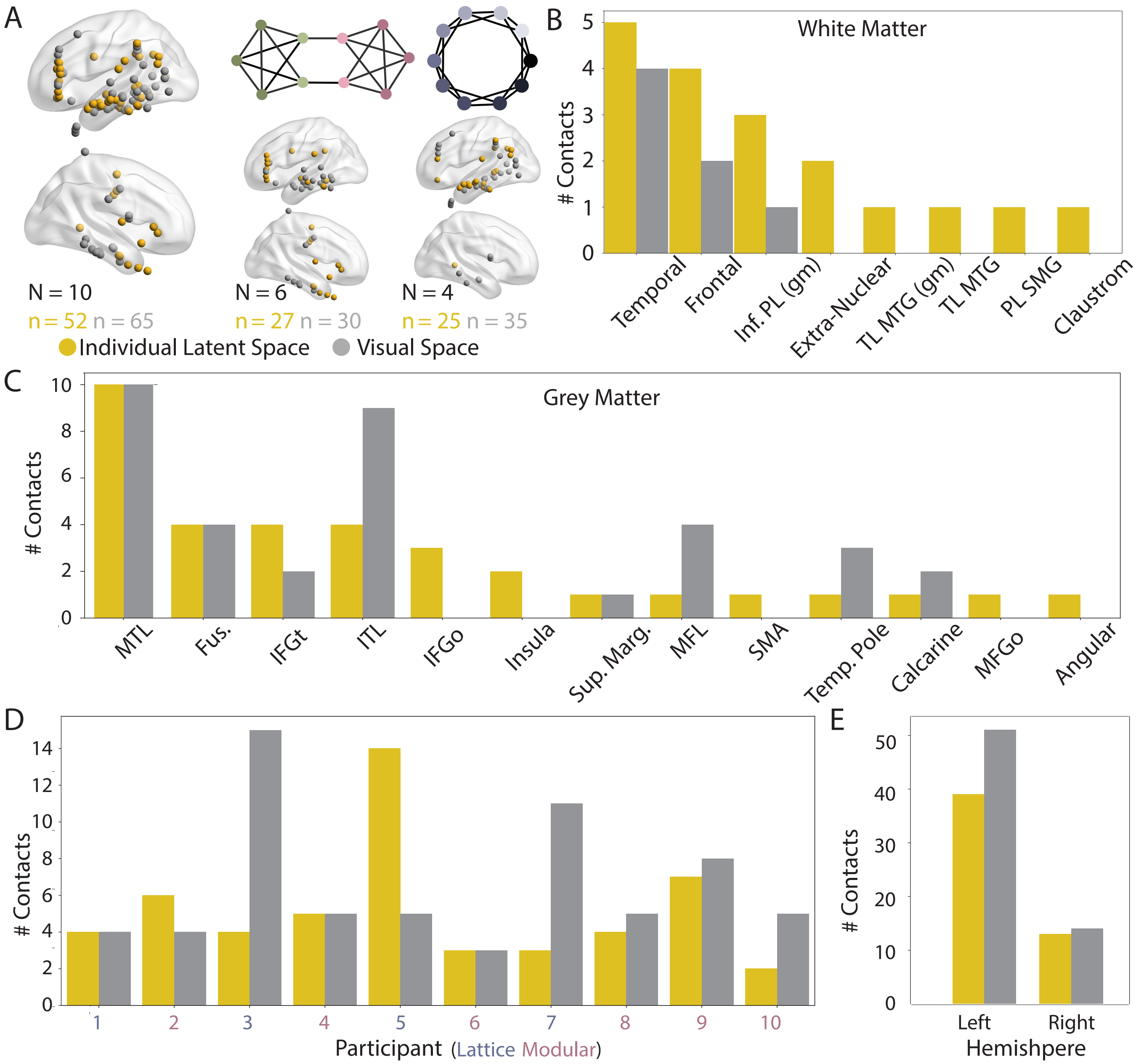


FIG. 5. **Middle-Aligned Contacts.** *(A)* Visualization of retained contacts from middle-aligned data with similar activity spaces on an MNI brain. Contacts that are similar to the latent space are shown in gold, and contacts that are similar to the visual space are shown in grey. *(B-C)* Anatomical localization of grey (*(B)*) and white (*(C)*) matter contacts. Grey matter contacts are localized with the AAL atlas, and white matter contacts are localized with the Talaraich atlas. *(D)* The number of contacts retained in each hemisphere for each space. *(E)* The number of contacts retained for each individual for each space. ITL: inferior temporal lobe, MTL: middle temporal lobe, TL MTG (gm): temporal lobe middle temporal gyrus (grey matter), Fus: fusiform, IFGt: inferior frontal gyrus (pars triangularis), IFGo: inferior frontal gyrus (pars orbitalis), MFL: middle frontal gyrus, SMA: supplemental motor area, Inf PL (gm): inferior parietal lobe (grey matter).

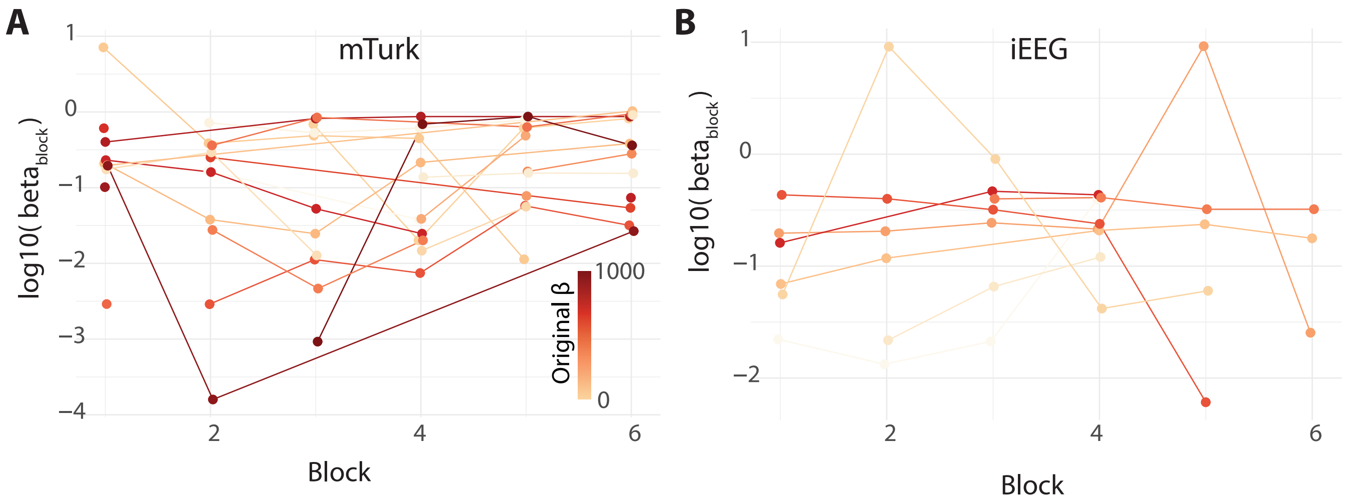


FIG. 6. **Time-varying Estimates of Beta.** *(A)* Beta estimates taken from 500 trial sliding windows of data from the mTurk cohort. The color of line is the $\beta$ value estimated from all the data. Missing values indicate the $\beta$ value was an extreme value of either 0 or 1000. *(B)* The same as in panel (A), but for the iEEG cohort.
